# Light-sheet microscopy with isotropic, sub-micron resolution and solvent-independent large-scale imaging

**DOI:** 10.1101/605493

**Authors:** Tonmoy Chakraborty, Meghan Driscoll, Malea Murphy, Philippe Roudot, Bo-Jui Chang, Saumya Vora, Wen Mai Wong, Cara Nielson, Hua Zhang, Vladimir Zhemkov, Chitkale Hiremath, Estanislao Daniel De La Cruz, Ilya Bezprozvanny, Hu Zhao, Raju Tomer, Rainer Heintzmann, Julian Meeks, Denise Marciano, Sean Morrison, Gaudenz Danuser, Kevin M. Dean, Reto Fiolka

## Abstract

We present cleared tissue Axially Swept Light-Sheet Microscopy (ctASLM), which achieves sub-micron isotropic resolution, high optical sectioning capability, and large field of view imaging (870×870 μm^2^) over a broad range of immersion media. ctASLM can image live, expanded, and both aqueous and organic chemically cleared tissue preparations and provides 2- to 5-fold better axial resolution than confocal or other reported cleared tissue light-sheet microscopes. We image millimeter-sized tissues with sub-micron 3D resolution, which enabled us to perform automated detection of cells and subcellular features such as dendritic spines.

---

Human tissues are composed of multiple polarized cell types organized in well-defined three-dimensional architectures. Interestingly, it has been shown that rare subsets of cells can play a crucial role in disease progression.^1^ Consequently, interdisciplinary efforts now aim to generate comprehensive atlases of human cells in diverse tissue types. To date, this has largely relied on massively parallel sequencing and machine learning-based analyses to identify distinct sub-populations of cells.^2^ Combined with advanced imaging, such efforts could not only shed light on the diversity of cell types, but the biological context in which each population operates. However, imaging large tissue with single cell and even subcellular resolution remains challenging because of the heterogeneous refractive index and composition of tissues, which results in complex aberrations and an increased scattering coefficient, both of which decrease spatial resolution and limit imaging depth.^3^

To circumvent this challenge, a large variety of optical clearing techniques have been developed that aim to homogenize the optical properties of the tissue using either aqueous^4–6^ or organic^7–9^ solvents. Today, these techniques now routinely render biological specimens sufficiently transparent, such that – if the imaging technology were to exist – entire organs could be imaged with subcellular resolution and molecular specificity.^10^ As the specimens are three-dimensional (3D), an ideal imaging system should possess isotropic, subcellular resolution, in order to accurately measure 3D cellular morphology and signaling activity.^11^ Nevertheless, imaging chemically cleared specimens with diffraction-limited or super-resolution presents technical challenges.^12^ For example, each clearing mechanism provides distinct advantages and disadvantages, and requires unique immersion media with refractive indices that range between 1.33 and 1.56. Thus, the imaging system must optimally operate throughout this refractive index range without suffering from deleterious aberrations, which decreases both resolution and sensitivity.

Light-sheet microscopy, because of its parallelized image acquisition, inherent optical sectioning, and ability to image large biological structures quickly and with high optical resolution, serves as an ideal candidate for cleared tissue imaging.^13, 14^ Nevertheless, to the best of our knowledge, there is not yet a light-sheet microscope for cleared tissue specimens that possesses submicron, isotropic resolution, and is also compatible with the full range of clearing methods.^7, 15–17^ Here, we address these limitations, and present a scalable imaging platform that provides sub-cellular anatomical detail in any spatial dimension across millimeter cubes of tissue. The system, which we refer to as cleared tissue Axially Swept Light-Sheet Microscopy (ctASLM), combines high-speed aberration-free remote focusing (**Supporting Note 1**), refractive index-independent illumination and detection optics, residual aberration correction, and syncronous camera readout to achieve sub-micron isotropic resolution throughout a volume of 870×870×5,000 μm^3^ with high optical sectioning capability (Figure 1a).^18, 19^ By tiling and stitching multiple volumes,^20^ ctASLM permits routine visualization of sub-cellular features throughout millimeters of tissue, regardless of the clearing method.

**Figure 1.**
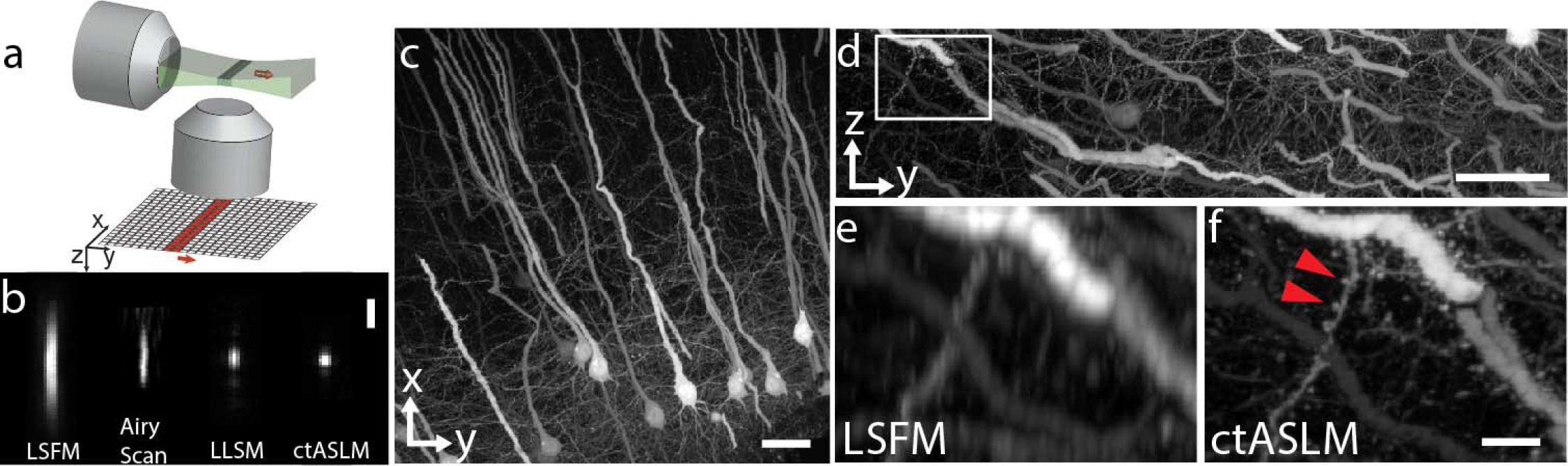
ctASLM enables isotropic, sub-micron imaging over large field of views. (a) Working principle of axially swept light-sheet microscopy: a thin light-sheet is scanned in its propagation direction, the rolling shutter readout of an sCMOS camera, adjusted to the size of the beam waist, is tightly synchronized to the light sheet scan. (b) Experimentally measured PSF of conventional light-sheet fluorescence microscopy (LSFM), Airy scan confocal microscopy, Lattice light sheet microscopy (LLSM) and ctASLM. The LLSM was using the same detection objective as the ctASLM system. (c-d) Maximum intensity projections of CLARITY cleared cortical Thy1-eYFP neurons as imaged by ctASLM. Zoomed in view of the rectangular region in (d) when imaged using, (e) LSFM vs. (f) ctASLM. All images show Raw data, no deconvolution was applied, with the exception of the Airy Scan PSF. Scale bars are: (b) 3 microns, (c,d) 100 microns; (f) 20 microns.

To evaluate the performance of the microscope, we imaged sub-diffraction beads with ctASLM, a conventional light-sheet microscope, an Airy Scan confocal microscope, and a lattice light-sheet microscope, each equipped with long working distance multi-immersion objectives (See Methods). Owing to the tradeoff between axial resolution and field of view (here 870 × 870 μm^2^), the conventional light-sheet microscope achieved a resolution of 1.04 ± 0.04 laterally and 6.7 ± 0.4 μm axially, respectively. Airy Scan improves both the lateral and axial resolution to 0.32 ± 0.03 and 3.1 ± 0.4 μm, respectively. Lattice light-sheet microscopy, adapted to cleared tissue imaging, improved the axial resolution to 1.53 ± 0.06 μm with a lateral resolution of 0.9 ± 0.1 μm, but at a cost of a 95-fold smaller field of view compared to the conventional light-sheet microscope.^21, 22^ In contrast, ctASLM achieved 0.90 ± 0.03 and 0.85 ± 0.03 μm lateral and axial resolution, respectively, over the full field of view (Figure 1b, Supplementary Figures 1 and 2, Supplementary Table 1, **and Supplementary Note 2**). Richardson-Lucy deconvolution further improved the resolution to 0.66 ± 0.03 lateral and 0.76 ± 0.02 μm axially (Supplementary Figure 3). Owing to ctASLM’s excellent optical sectioning and high-quality raw data, deconvolution is not mandatory (Supplementary Figure 4). Indeed, neurons could be traced in any spatial dimension simply using the unprocessed data (Figure 1c, **and** 1d). In comparison to a conventional light-sheet microscope (Figure 1e), individual synaptic spines were clearly resolved even in the axial direction (Figure 1f).

Importantly, different biological tissues require specific clearing methods, each with their own solvents and labeling strategies. Thus, we sought to evaluate ctASLM’s performance in solvents with refractive indices between 1.333 (water) and 1.559 (BABB). In aqueous solutions, we resolved fine vascular detail in a live zebrafish expressing an endothelial reporter (Figure 2a), and visualized large-scale dendritic projections in chemically expanded hippocampal slices (Supporting Figure 5, **and** Supporting Movie 1). In the intermediate refractive index range of ~1.44, we imaged a portion of a CLARITY-cleared mouse brain spanning 5×3×3 mm^3^, and the high spatial resolution permitted a 3D segmentation of individual neurons in densely packed cortical regions (Figure 2b **and** c, Supporting Movie 2 **and** 3). Using the chemical clearing method PEGASOS, we imaged the entire murine olfactory bulb (Supporting Movie 4). Lastly, we imaged an entire BABB cleared neonatal kidney, labeled with a vasculature marker Flk1-GFP, encompassing 3.4×2.6×2.5 mm^3^ (Figure 2d, Supporting Movie 5). Throughout the entire volume, we maintained sufficiently high spatial resolution to distinguish individual nuclei (Figure 2e, Supplementary Figure 6).

**Figure 2.**
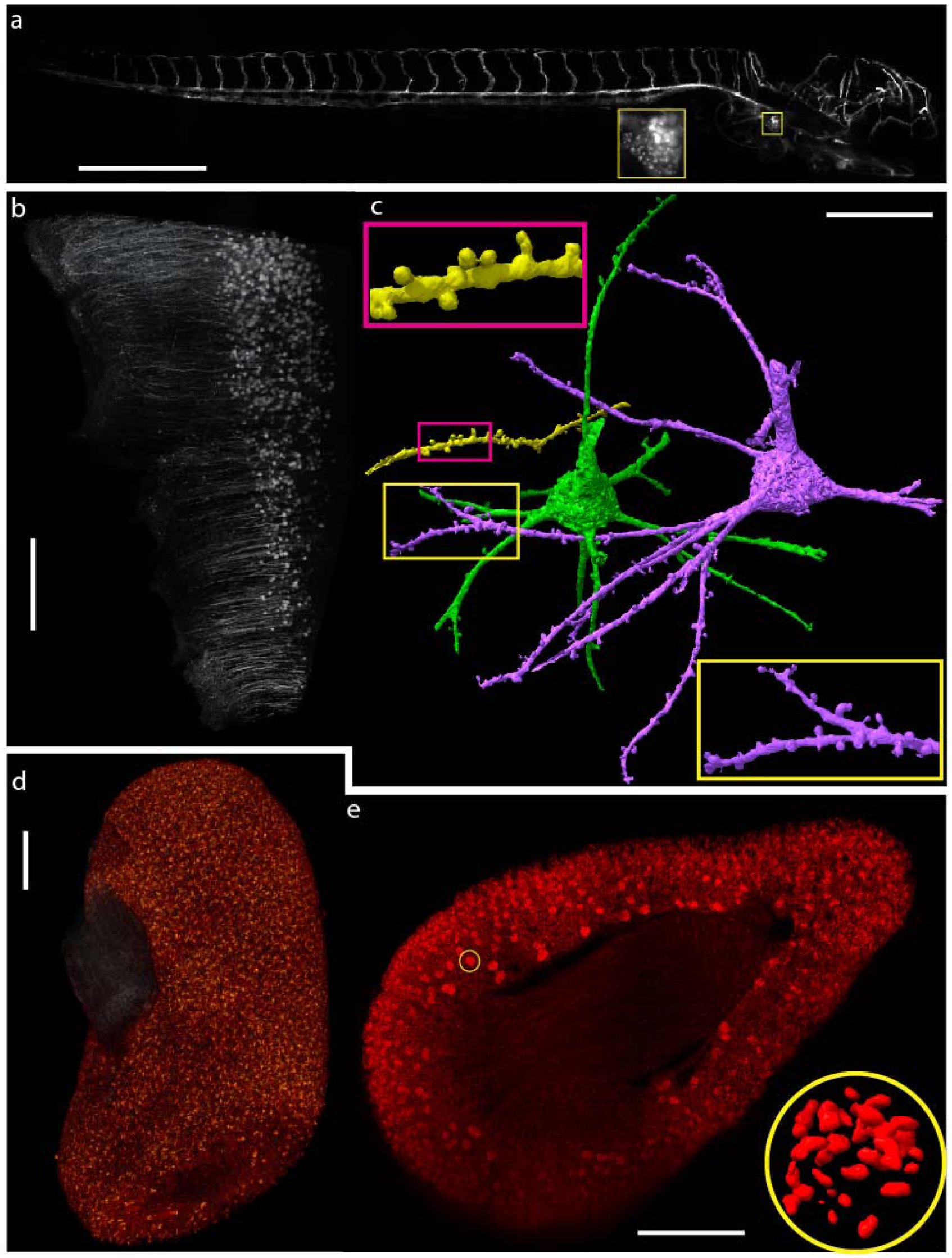
ctASLM allows isotropic, submicron imaging over a wide refractive index range. (a) Vasculature of a live 72-hour old Zebra fish imaged in E3 medium. Vasculature endothelial cells labeled with mCherry. The inset shows a zoomed in view of the heart. Scale bar, 500 μm. (b) Thy1-eYFP neurons in a mouse brain cleared with CLARITY. Scale bar, 500 μm. (c) Segmentation of two neurons (magenta and green) and a dendrite from a neighboring neuron (yellow) from the dataset shown in (a), Scale bar, 50 μm. Insets shows magnified views of dendrites and their spines. (d) Volume rendering of a mouse kidney labeled with Flk1-GFP. Scale bar, 500 μm. (e) one cross-section through the kidney shown in (d), Scale bar, 500 μm. Inset shows a 3D rendering of endothelial cells in one selected Glomeruli.

Given ctASLM’s isotropic resolution, large field of view imaging, and excellent optical sectioning, we were able to extract quantitative biological information from our imaging data. Using open source software, we were able to not only readily detect dendritic spines but also cluster spines based on their 3D morphology (Figure 3a, **and** Supplementary Movies 6 **and** 7).^23^ These clusters span a wide range of spine morphologies (Figure 3b), form a structured space following principal component analysis (Figure 3c), and are separable via interpretable measures, such as the ratio of spine neck area to spine surface area (Supplementary Figure 7). For the kidney, glomeruli serve as the most basic filtration unit, and damage to glomeruli is associated with a spectrum of clinical outcomes, including proteinuria and reduced filtration, which are the hallmarks of chronic kidney diseases. To evaluate tissue composition in an automated fashion, we developed a multiscale watershed algorithm to identify individual glomeruli (Figure 3d). Further, owing to the high axial resolution, we can detect endothelial cells within an individual glomerulus (Figure 3e, Supplementary Figure 8, Supporting Movie 8). Formerly, such an analysis was performed manually on hundreds of serial-sectioned tissue preparations, as identification of individual cells in confocal datasets was particularly challenging due to its diminished z-resolution.

**Figure 3.**
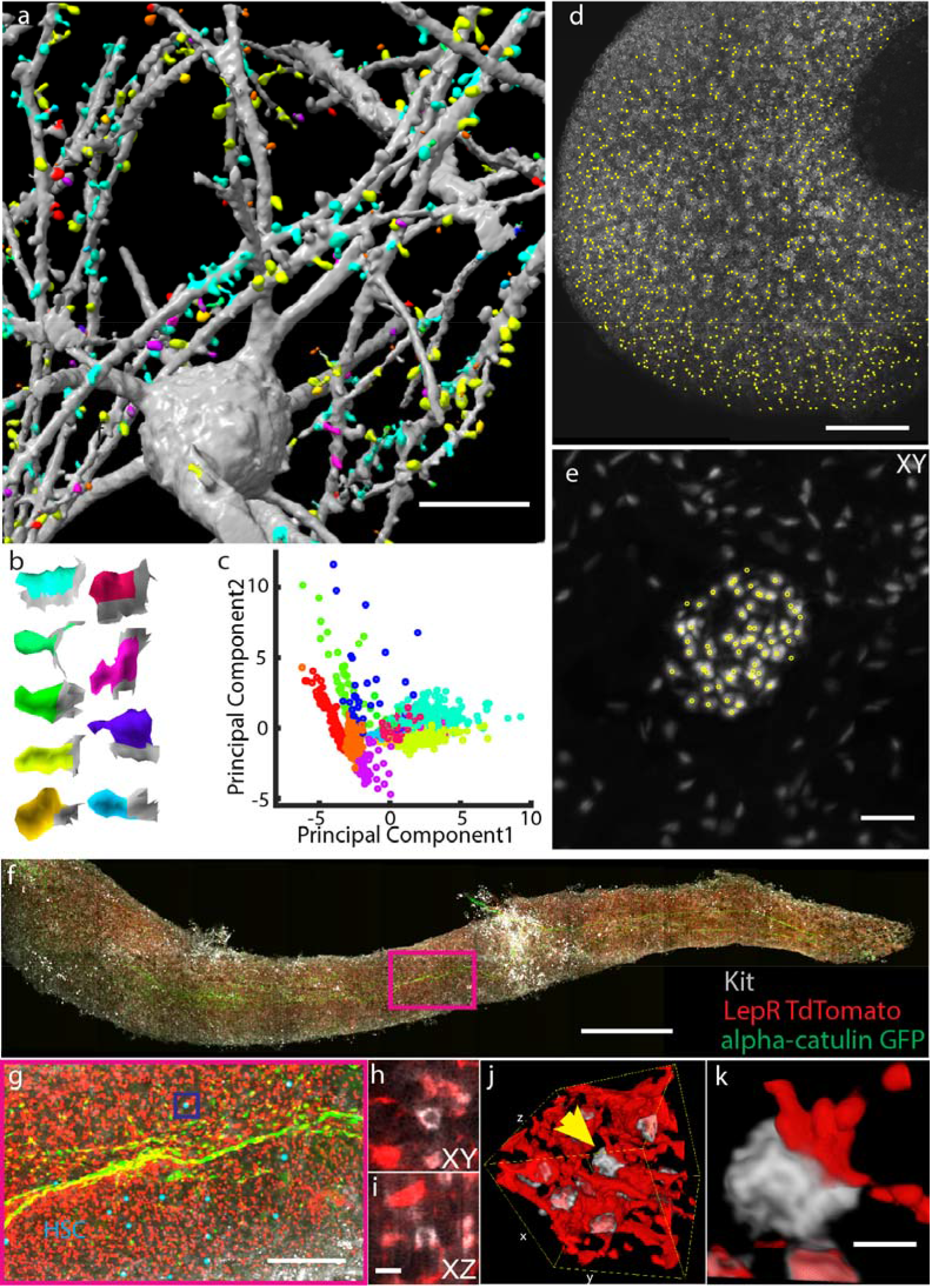
ctASLM allows detection of cellular and subcellular features in large tissues. (a) Machine learning based supervised detection and unsupervised clustering of spines. (b) The shape of the median spine in each cluster. (c) Principal component analysis of the spines. (d) automatic detection of glomeruli in the mouse kidney data set shown in Fig 2 (d). (e) Lateral maximum intensity projection of one Glomeruli and detection of individual endothelial cells (yellow circles). (f) Maximum intensity projection of a mouse tibia bone marrow plug. (g) Zoom in of magenta box shows Apha-catulin-GFP+(green), Kit+(grey) HSCs labeled with cyan spheres. (h-l) cross-sectional views of Kit+ HSCs and LepR TdTomato+ niche cells (red). (h&i) are the raw data and (j) is the volume-rendering of boxed region in (g).(k) Scale bar: (a) 20μm (d) 200μm (e) 50μm (f) 1mm (g) 200μm (h-i) 10 μm (j) 40×40×40 ◻m cube (k) 5 μm

Lastly, we imaged hematopoietic stem cells (HSCs) in mouse bone marrow. HSC are responsible for the continued production of blood and immune cells throughout life. HSCs reside within a perisinusoidal niche in the bone marrow where Leptin Receptor+ (LepR+) stromal cells and endothelial cells synthesize the factors required for HSC maintenance. Previous studies have shown that LepR+ cells and HSCs are in close proximity^24^. However, careful analysis of the physical interactions between these cells has not been done, as HSCs are a rare population that makes up only 0.003% of all hematopoietic cells within the bone marrow. Therefore, we imaged an entire bone marrow plug with ctASLM (Figure 3h), which allowed us to identify all a-catulin^+^c-kit^+^ HSCs, markers that identify all HSCs in young adult bone marrow and which give very high levels of purity (Figure 3i).^24^ Sub-micron isotropic resolution reveals extensive physical interactions of LepR+ cell projections and HSCs (Figure 3j-n, Supplementary Movie 9), which are distorted when imaged with the anisotropic resolution of an Airyscan confocal microscope.

In summary, ctASLM provides subcellular detail and tissue scale anatomy. As tissues are inherently 3D, the isotropic resolution of ctASLM is an important attribute to describe 3D morphologies and molecular concentrations in an unbiased way. ctASLM imposes a higher out-of-focus excitation load on the sample than a conventional light-sheet system and illuminates a smaller portion at a time, both of which can increase photobleaching. While we were not limited by photo-bleaching on the samples we imaged, we note that ctASLM is compatible with light-sheet engineering via Field Synthesis to optimize confinement and axial resolution (**Supplementary Note 3**). We anticipate that higher overall 3D resolution can be achieved by the development of higher NA, long working distance objectives. Such a system will likely require adaptive optics to realize the best resolution deep in tissues, a need we already see for our current system when imaging in CLARITY cleared brains (Supplementary Figure 6). With the given resolution level, we readily show that we can combine ctASLM with computer vision to detect and classify biological features throughout millimeters of tissue. Consequently, we believe that ctASLM will expedite human cell atlas efforts, providing much-needed insight into how tissue function manifests in both health and disease, from the heterogenous cellular populations that compose it.

## Acknowledgements

We would like to thank the Cancer Prevention Research Institute of Texas (RR160057 to R.F., and R1225 to G.D.) for their generous funding, as well as the National Institutes of Health (F32GM116370 and K99GM123221 to M.K.D., and R01GM067230 to G.D., R01AG055577 and R01NS056224 to I.B., R01DK118032 and R01DK099478 to M.B., R01DC015784 and R21NS104826 to J.B. and R33CA235254 to R.F.). We are grateful to the Live cell imaging facility at UT Southwestern for access to the Zeiss Airy Scan confocal microscope (which is supported by 1S10OD021684-01 to Katherine Luby-Phelps). We would like to thank David Saucier and Dr. James Amatruda for providing the zebrafish specimens, and Dr. Etai Sapoznik for his assistance with CAD design.

## Author Contributions

T.C., K.M.D., and R.F. designed the research. T.C. and R.F designed and T.C. built the microscope. T.C., and S.V. prepared the CAD files and operated the microscope. T.C., M.K.D., K.M.D., B-J.C, P.R., and R.F. performed image analysis, partially under guidance by G.D. R.H. simulated PSFs. W.M.W., C.N., H.Z., V.Z., M.M., C.H., D.M., I.B., H.Z., R.T., J.M., and S.M. provided specimens and guided imaging. T.C., K.M.D., and R.F. wrote the manuscript. All authors read and provided feedback on the final manuscript.

## Additional Information

The datasets acquired for this study are available from the corresponding author upon reasonable request.

## Supplementary Methods

### Animal Specimens

All animal protocols were approved by local Institutional Animal Care and Use Committees (IACUC) as directed by the National Instutes of Health, and strictly followed. These protocols include AC-AAAR0417 (to R.T., Columbia University), 2017-102370 (to J.M., UT Southwestern Medical Center), 101917 (to D.M., UT Southwestern Medical Center), 101715 (to I.B., UT Southwestern Medical Center), and 102632 (to S.M., UT Southwestern Medical Center).

### Microscope Control

The data acquisition computer was a Dell Precision 5810 Tower equipped with an Intel Xeon E5-1650W v3 processor operating at 3.5 GHz with 6 cores and 12 threads, 128 GB of 2133 MHz DDR4 RAM, and an integrated Intel AHCI chipset controlling 4× 512 GB SSDs in a RAID0 configuration. All software was developed using a 64-bit version of LabView 2016 equipped with the LabView Run-Time Engine, Vision Development Module, Vision Run-Time Module and all appropriate device drivers, including NI-RIO Drivers (National Instruments). Software communicated with the camera (Flash 4.0, Hamamatsu) via the DCAM-API for the Active Silicon Firebird frame-grabber and delivered a series of deterministic TTL triggers with a field programmable gate array (PCIe 7852R, National Instruments). These triggers included control of the resonant mirror galvanometers, voice coils, stage positioning, laser modulation and blanking, camera fire and external trigger. Some of the core functions and routines in the microscope control software are licensed under an MTA from Howard Hughes Medical Institute, Janelia Farm Research Campus. The control software code for ctASLM can be requested from the corresponding authors and will be distributed under MTA with HHMI Janelia Research Campus.

### Microscope Layout

A schematic layout of the ctASLM microscope is shown in Supplementary Figure 9 and a part list is provided in Supplementary Table 2. The illumination train consisted of four Coherent Obis lasers (LX 405-100C, LX 488-50C, LS 561-50, LX 637-140C) that were combined with dichroic beam splitters (LM01-427-25, LM01-503-25, LM01-613-25, Semrock), focused through a 30 μm micron pinhole (P30D, ThorLabs), with a 50 mm achromatic doublet (AC254-50-A, ThorLabs), recollimated with a 200 mm achromatic doublet, (AC254-200-A-ML, ThorLabs), and directed through a 3× Galilean beam expander (GEB03-A). Thus, the initial beams were expanded by a factor of 12× before being focused onto a resonant mirror galvanometer (CRS 4 kHz, Cambridge Technology) with a cylindrical lens (ACY254-50-A, ThorLabs). The mirror galvanometer was used to wobble the light sheet and was driven with a 12V power supply (A12MT400, Acopian), and the 1D focus of the cylindrical lens was relayed with a 200 mm achromatic doublet (AC508-200-A, ThorLabs), passed through a 50:50 polarizing beam splitter and a quarter waveplate, and imaged onto a mirror with a 4× microscope objective (XL Fluor 4×/340, NA 0.28, Olympus Life Sciences). The mirror was mounted on a voice coil with 10 mm of travel, sub-50 nm positioning repeatability, and a sub-3 millisecond response time (LFA-2010, Equipment Solutions). The reflected light was recollected by the same 4× objective, the polarization state was rotated by the second passage through the quarter waveplate (AQWP3, Boldervision) reflected with the cube beam splitter (10FC16PB.7, Newport) and relayed to the illumination objective (Cleared Tissue Objective, NA 0.4 Advanced Scientific Imaging) with 200 mm and 75 mm achromatic doublets (AC508-200-A and AC508-75-A, ThorLabs). To correct for spherical aberrations, 3 mm N-BK7 glass piece (37-005, Edmund Optics) was placed between the 4× objective and the voice coil actuated mirror (Supplementary Figure 10).

The detection arm consisted of an identical microscope objective (Cleared Tissue Objective, Advanced Scientific Imaging), a tube lens (ITL200-A, ThorLabs), a 4-position filter wheel (LB10-WHS4 and a Lambda 10-3, Sutter Instruments), and a sCMOS camera (Flash 4.0, Hamamatsu Corporation). For emission filters, we used 445/20, 525/30, 605/15, and 647 long-pass filters (FF01-444/20-25, FF01-525/30-25, FF01-505/15-25, and BLP01-647R-25, Semrock), for blue, green, red, and far-red, respectively. Both the illumination and detection objectives were immersed in a large chamber that was used to house the specimen. To collect a Z-stack, a voice coil direct drive was used to scan the specimen in a sample-scanning mode (V-522.1AA, C-413.2GA, and C-413.1IO, Physik Instrumente). Sub-volumes were acquired by moving the position of the specimen and the voice coil stage with a 3-axis motorized stage (3DMS and MP-285A, Sutter Instruments). To image a 1mm^3^ volume using ctASLM takes 17.6 minutes, using 100 ms exposure time, an isotropic step size of 0.425 μm and 18% overlap between stacks for faithful stitching.

### Sample mounting

Samples were glued with cyanoacrylate (PT09, Pacer Technology, California) or with silicone (GE Silicone 2+) on to a 1×2.5 cm^2^ glass slide cut from a 2.5×7.5 cm^2^ microscope slide (Cat. No. 3050, Thermo Scientific). The glass slides were mounted with a custom holder to the sample scanning stage. CAD drawings of the sample holder assembly are shown in Supplementary Figure 11.

### Low-NA Light-Sheet Microscopy

For low-NA light-sheet imaging, the same optical train was used as in ctASLM, but with the presence of a variable width slit aperture conjugate to the back pupil of the cylindrical lens, and without axial scanning of the illumination beam. The illumination NA was selected such that the entire field of view (870 × 870 μm^2^) resided within two Rayleigh lengths of the illumination beam. This resulted in a Gaussian beam with a e^−2^ thickness of ~7 μm, which is in agreement with resolution measurements obtained using sub-diffraction fluorescence beads.

### Lattice Light-Sheet Microscopy

Current lattice light-sheet microscopes are designed to operate with high numerical aperture objectives optimized for water dipping only. To compare the performance of LLSM with ctASLM in terms of multi-immersion imaging, we modified an existing lattice light-sheet setup^25^ to measure the effects on the PSF when the lower NA cleared tissue objectives are used in LLSM. To this end, we mounted one Cleared Tissue Objective (NA 0.4, Advanced Scientific Imaging) in the detection path of the LLSM. On the illumination path, we used a NA 0.8 40× (Nikon) objective, for which the rest of the illumination train was optimized. However, we limited the used NA to less than 0.4 to simulate the lower NA available if a cleared tissue objective is used. We did not employ directly a cleared tissue objective in the illumination path, as this would have required a complete redesign of the optical train in the illumination path. We used the square lattice mode, as this is the most widely used lattice pattern, also in the most recent LLSM imaging of expanded tissues. We imaged 200nm green fluorescent microspheres in agarose and water immersion. We estimated axial resolution and the illuminated field of view from the bead images.

### Confocal Microscopy

In order to compare the performance of the ctASLM to that of the commercial microscopes, we compared the z-resolution performance of Airyscan and a confocal microscope to our ctASLM. Imaging of the bone plug was performed on a Zeiss LSM880 equipped with a long working-distance multi-immersion objective (LD LCI Plan-Apo 25× 0.8 NA) as previously described.^24, 26^ PEGASOS cleared mouse brain were also imaged using a Leica SP8 confocal laser scanning microscope equipped with an HCX APO L20x/0.95 immersion objective. For best performance a 56μm pinhole size with a pixel dwell time of 1.84us was used. A comparison of ctASLM to the data acquired with the Leica SP8 is shown in Supplementary Figure 12.

### Sample Preparation

#### Zebrafish

The zebrafish embryo is a transgenic line (krdl:mCherry) expressing mCherry on the vascular endothelium. A 3-day post fertilization zebrafish embryo is mounted in 2% low-melting agarose (A9045-25G, Sigma-Aldrich) in an FEP (Fluorinated ethylene propylene) tube (FEP HS .029 EXP/ .018 REC, Zeus). The working solvent of the agarose is E3 fish water. The FEP tube is sonicated in 70% Ethanol and then washed with clean distilled water prior to the mounting of the zebrafish embryo. We first pick and leave a zebrafish embryo in a few (~50 μL) E3 media on a petri dish. Next, 5 μL of 0.1% Tricaine is added to anesthetize the zebrafish embryo. About 50 μL of the mixture of the Tricaine and E3 medium is removed, and then the liquid low-melting agarose is added on the zebrafish embryo. The zebrafish embryo in the liquid agarose is aspirated directly into the clean FEP tube with a pipette. Within a few minutes, the agarose is solidified, and the zebrafish embryo is mounted in the FEP tube. For the last step, we use a needle to poke two holes on the FEP tube above and below the zebrafish embryo to increase the gaseous exchange allowing the embryo to breathe. The zebrafish embryo is then transferred into the microscope chamber and ready to be imaged.

#### CLARITY Brain

*Thy1-eYFP* brain were extracted and cleared using passive CLARITY^27^ with the following hydrogel monomer (HM) solution recipe: 1% (wt/vol) acrylamide, 0.05% (wt/vol) bisacrylamide, 4% paraformaldehyde, 1× Phosphate Buffered Saline, deionized water, and 0.25% thermal initiation VA-044 (Wako Chemicals, NC0632395). The transcardiac perfusion was performed with 20 ml HM solution, followed by brain extraction and overnight incubation at 4°C. The Hydrogel was polymerized by incubating the brain samples at 37°C for 3-4 hours, and the clearing was performed by shaking in clearing buffer (4% (wt/vol) SDS, 0.2 M boric acid, pH 8.5 at 37°C, ~3 weeks). After washing with the buffered solution (0.2 M boric acid buffer, pH 7.5, 0.1% Triton X-100), the brains were transferred into 50-87% glycerol solution for refractive index matching (as described previously^28^) and imaging.

#### Olfactory Bulb

Double transgenic mice were made by crossing Ai9 mice(B6.Cg-Gt(ROSA)26Sortm9(CAG-tdTomato)Hze/J, Jax Laboratory Stock No. 007909) with FosTRAP mice^29^ and resulting mice were termed cFosxAi9 mice. Briefly, animals were anesthetized with IP injections of ketamine/xylazine (120mg/kg ketamine and 16mg/kg xylazine). After verification of complete anesthesia via the toe pinch test, the chest cavity was exposed and a syringe needle was inserted into the left ventricle. Mice were perfused with 20mL of PBS solution followed by 20mL of a 4% PFA Solution. After perfusion, mice were decapitated and the brain, including the olfactory bulbs, were carefully dissected out and placed into 4% PFA solution overnight. These samples were then cleared using the PEGASOS method as described in Jing et al.^30^ All samples were kept at 42°C for this process. Briefly, samples were decolorized for 2 days in 25% Quadrol before undergoing delipidation in a series of 30%, 50%, and 75% tert-Butanol solutions. This was followed by a final delipidation step with TBP (75% tert-butanol and 25% PEGMMA solution). Finally, samples were submerged in BB-PEG (75% benzyl benzoate, 22% PEGMMA, 3% Quadrol) until imaging. These data shown here were based upon a male mouse aged 3.32 months.

#### Bone Marrow

Processing of these samples were carried out as previously described in Acar et al.^24^ Briefly, intact bone marrow plugs from freshly dissected tibias of 6–8-week-old female mice were extruded from the bone using a PFA-filled syringe with a 25-gauge needle and placed directly into 4% PFA solution for 6 h at room temperature. Fixed plugs were then washed in PBS and blocked in whole mount staining medium (PBS with 5% donkey serum, 0.5% NP-40, and 10% DMSO) overnight at room temperature. After blocking, plugs were stained for three days at room temperature with primary antibodies (chicken anti-GFP: GFP-1020, Aves Labs and goat anti-c-kit: BAF1356, R&D Systems) in staining solution. Then the tissues were washed multiple times in PBS at room temperature for one day and put into staining solution containing secondary antibodies (Alexa Fluor 488-AffiniPure F(ab')2 fragment donkey anti-chicken IgY and Alexa Fluor 647-AffiniPure F(ab')2 fragment donkey anti-goat IgG) for three days followed by a one-day wash to remove any unbound secondary antibodies. Samples were cleared using PEGASOS clearing protocol, previously described in Jing et al.^30^ Briefly, marrow plugs were embedded in 2% low melt agarose in water and were dehydrated in a series of 30%, 50%, 70%, and 100% Tert-Butanol. Each step was performed for 8 hours at room temperature. Samples were then refractive index matched overnight at room temperature in BB-PEG. BB-PEG was prepared as a 3:1 mixture of benzyl benzoate (BB) (Sigma-Aldrich B6630) and PEGMMA500 (Sigma-Aldrich 447943) supplemented with 3% w/v Quadrol (Sigma-Aldrich 122262). Samples were imaged 1 day to 2 months after initial refractive index matching without significant loss of endogenous TdTomato or Alexa Fluor signal.

#### Isolated Hippocampus

Thy1-GFP mouse was transcardially perfused with cold PBS first, then 4% PFA in PBS for fixation. The brain was taken out and put in 4% PFA in PBS overnight at 4°C, then 2 mm thickness hippocampal slices was cut with vibratome coronally. The hippocampal slices were then cleared with PEGASOS clearing protocol, previously described in Jing et al.^30^ Briefly, the slices were decolorized in 25% Quadrol for 1 day, then delipidated in series of 30%, 50%, 70% Tert-Butanol solutions, dehydrated in tB-PEG, and finally cleared in BB-PEG.

#### Expanded Hippocampus

Thy1-GFP mouse was transcardially perfused with cold PBS first, then 4% PFA in PBS for fixation. The brain was taken out and put in 4% PFA in PBS overnight at 4°C, then 0.1 mm thickness hippocampal slices were cut with vibratome coronally. Samples were processed according to the original protein-expansion microscopy procedure.^21, 31^ Briefly, slices were permeabilized using 5% BSA, 0.5% Triton X-100 buffer for 1 hour and incubated overnight with a coupling reagent 0.1% acryoyl-X succinimidyl ester in PBS.^31^ Slices were washed 3 times with PBS for 5 minutes and incubated in polymerization solution (8.6% sodium acrylate, 2.5%, 0.15% N,N – Methylenebisacrylamide, 2 M NaCl, 0.2% APS, 0.2% TEMED, 0.01% TEMPO) for 1 h at 4 C before prior to polymerization at 37 C for 2 h. Embedded tissue was transferred to isotonic proteinase K buffer (50 mM Tris-HCl pH 8.0, 2 M NaCl, 1 mM EDTA, 0.5% Triton X-100, proK 8 u/mL) and incubated for 2 h at room temperature. Complete proK digestion was achieved overnight at room temperature in proteinase K buffer (50 mM Tris-HCl pH 8.0, 1 M NaCl, 1 mM EDTA, 0.5% Triton X-100, proK 8 u/mL)1. Next day, samples were expanded for 1 h in double-distilled H_2_O with 3 water changes. Glass holders were covered with 0.1% poly-L-lysine for 1 h, washed 3 times with ddH_2_O and allowed to air dry for 1 h. Hippocampus region was excised from coronary slices and mounted on a poly-L-lysine coated glass holder. For long-term storage and image acquisition samples were transferred to 1 mM NaOH buffered solution.

#### Neonatal Kidney Preparation

Post-natal day 3 kidneys from an Flk1-GFP (Jackson Laboratories, 017006) mouse were harvested and fixed for 2h at room temperature in 4% paraformaldehyde in PBS. Procedures were performed according to UTSW-IACUC-approved guidelines. Fixed kidneys were incubated with tert-Butanol/water mixtures of increasing alcohol concentrations (30%, 50%, 70%, 80%, 96%, and twice 100%) with 2% Triethanolamine. The kidneys were then left in BABB (benzyl alcohol and benzyl benzoate in a 1: 2 volume ratio) with 10% Triethanolamine until tissue cleared. For prolonged storage in clearing medium (> 2 days), samples were stored at 4°C.

### Deconvolution, stitching, and 3D visualization

Image processing was performed on a Windows 10-based workstation equipped with two Intel Xeon Gold 5120 CPUs, 1 TB of RAM, an NVIDIA Quadro P6000 GPU. The point spread function (PSF) used for deconvolution was synthesized from the imaging parameters (wavelength, numerical aperture, refractive index) and 3D-deconvolution was performed with 40 iterations of the Richardson-Lucy algorithm (Microvolution). To stich the sub-volumes, the Fiji-based plugin BigStitcher^32^ was used. Image analysis was performed with Fiji^33^ and MATLAB (Mathworks), and 3D renderings were produced with ChimeraX or Arivis.

### Data analysis

#### HSC Detection

HSCs were detected as previously described.^24^ Briefly, individual alpha-catulin-GFP+, Kit+ HSCs were identified manually using the orthoslicer function of Bitplane Imaris v9.2.1 software. HSC coordinates and size were interactively annotated using the Imaris spots function in manual mode. Kit and LepR TdTomato surfaces were generated in Imaris using iso-surface function based on an empirically chosen intensity threshold.

#### Glomeruli Detection

In an effort to automatically detect the abundance of glomeruli throughout the entire neonatal kidney tissue, sub-volumes (1024 × 1024 × 470 pixels) were retrieved from the stitched volume (3042 × 4018 × 2757 pixels) and loaded into MATLAB. Sub-volumes were subjected to a three-dimensional Gaussian blur of 20 pixels before individual glomeruli were detected using an auto-adaptive watershed algorithm implemented in MATLAB.^34^ This algorithm uses a difference of Gaussians to suppress features that are either too large or small, here defined as 100 and 10 pixels, respectively. Segmented objects touching the border of the sub-volume were removed. After identification of glomeruli, the encompassing region was again loaded into MATLAB, and the full resolution was subjected to the same watershed algorithm, but with a difference of Gaussian of 10 and 2 pixels, respectively.

#### Spine Detection and Clustering

We detected spines using previously described software.^23^ Images were first deconvolved for 10 iterations using the MATLAB function *deconvblind()*, which uses a maximum likelihood algorithm to infer the fluorescence distribution in the sample. Following a gamma transform of 0.4 and 3D hole filling, a triangle mesh representing the cell surface was created as an isosurface at the Otsu threshold value of the image. The mesh was next decomposed into approximately convex patches using the parameters previously described,^23^ except that merging via the triangle criterion was disabled and merging via the line-of-sight criterion was performed more conservatively with a parameter of 0.8. Training data for patch merging and patch classification machine-learning models were then generated from nine non-overlapping images of size 200 pixels in the lateral dimensions and approximately 150 pixels in the axial dimension. These nine images were cropped from eight larger images. Using leave-one-out cross validation on patches labeled as certainly spines or certainly not spines, we calculated that our spine detection workflow had a precision of 0.86 and a recall of 0.88.

We performed an unsupervised clustering of detected spines into 15 clusters using agglomerative hierarchical clustering. For each cluster, we calculated the median of each of the morphological feature vectors used for clustering and plotted the spine closest to that median (Figure 3b). Only the clusters with a median spine attached to the neuron body were plotted. Principal component analysis was performed on these same features. We defined the spine neck surface area, as the patch closure surface area. We previously defined^23^ the closure surface area as the additional surface area needed to minimally generate a closed polyhedron from the open polyhedron comprising the dendritic spine. We also defined the variation from a sphere by fitting each spine to a sphere and measuring the ratio of the standard deviation of distances from the sphere at each mesh face to the sphere radius.

#### Segmentation of Neurons

The mouse brain volume was tiled in sub-volumes for parallel processing, and the signal was enhanced using a volumetric Laplacian-of-Gaussian transform with an isotropic scale of 1.5 voxels. The resulting volume was then transformed into a triangular mesh using an empirically chosen threshold. In order to reduce clutter, only the connected components with volume superior to the 99 percentiles of all connected component volume are displayed in Figure 2c. The algorithm is implemented using the open-source G’MIC software (https://gmic.eu), and the mesh was imported into Meshlab^35^ for coloring and image capture.

#### Statistics and Reproducibility

For statistical analysis, we report the mean, standard deviation, and number of observations. For PSF measurements of ctASLM, lattice light-sheet and conventional light-sheet, the full width half maximum (FWHM) for five green fluorescent beads was measured. For the Airy Scan microscope and the deconvolved ctASLM data, the FWHM for three beads was measured. Zebrafish embryos were imaged three times. Only one CLARITY brain sample was used, but it was imaged multiple times. Two expanded brain tissue slices were produced and imaged. Segmentation of Neurons was perfomed on one data set from a CLARITY brain sub-volume and on one data set from a hippocampal slice sub-volume with PEGASOS. In all cases image performance was checked by measuring point or filament like objects and image blur was analyzed by looking at cross-sectional slices. Synaptic spines were detected on two different sub-volumes.

### Data Availability

The datasets acquired for this study are available from the corresponding author upon request.

### Code Availability

The instrument control software can be requested for academic use from the corresponding authors and will be delivered under material transfer agreements with HHMI and UT Southwestern Medical Center.

### Supplementary Notes

#### Note 1 – Remote focusing technologies for ASLM

Various actuation principles have been used to scan light-sheets in their propagation direction, including resonant acoustic lenses,^36^ binary holograms,^22, 37^ electro-tunable lenses,^38^ and aberration-free remote focusing.^19^ While most of these techniques simply introduce a defocus term to the wavefront, only remote focusing with a matched secondary objective compensates for higher order aberrations. Simulations assuming ideal operations of an electro-tunable lens showed that the Strehl ratio would drop by 28%, and that the confinement of the light-sheet gets worse. For these reasons we chose remote focusing with a separate objective. Furthermore, the multi-immersion objectives do not possess a correction collar, however, remote focusing allows us to compensate for spherical aberration: by using a remote air objective that was originally designed to image through a small layer of water, this opens the possibility for remote aberration correction: We can compensate remaining spherical aberrations (imperfection of lenses, but also chromatically varying aberrations) of the system by inserting or removing blocks of glass. Note that with a purely air corrected objective (coverslip-free design), one could only correct one sign of spherical aberration, as one can only insert materials with a refractive index higher than one. With the objective designed for partial imaging in air and water, we can correct in principle remotely for spherical aberrations of either sign. Another requirement for ASLM using medium to high NA objectives is the precise waveform control. We found that using the voice coil actuator, we could achieve sufficiently linear remote focusing scans at 10 Hz. This corresponds to a maximum imaging frame rate of 100 ms, which we found in practice to be a suitable imaging rate for most samples (balance between acquisition rate, signal strength and needed excitation power).

#### Note 2 – Axial resolution and multi-immersion compatibility

In an epifluorescence microscope, axial resolution is determined by the numerical aperture of the objective. For cleared tissue imaging, the highest NA objective optimized for visible wavelengths is to our knowledge the HCX APO L20x/0.95 immersion objective (Leica). However, even using this objective for confocal microscopy, the axial resolution is ~3 microns, which is insufficient to resolve subcellular features such as spines (Supplementary Figure 12). For light-sheet microscopy, the axial resolution is governed by the product of the axial excitation intensity profile and the detection PSF. For conventional Gaussian light-sheets, the width of the sheet may exceed the axial extent of the detection PSF. In this scenario, axial resolution is mainly dominated by the detection NA, which explains the rather poor axial resolution we measured on our conventional light-sheet implementation.

Given that ctASLM is implemented with two identical objectives for excitation and detection, and the full NA is used for light-sheet generation, the axial and lateral resolution become close to identical. As the width of the illumination beam is narrower than the extent of the depth of focus (~6 microns), the axial resolution is mainly determined by how thin the light-sheet is (e.g., the illumination NA). This property allows ctASLM to use recently developed multi-immersion objectives of medium NA (0.4) for both detection and excitation. These objectives were designed to operate without a correction collar, but instead feature a concave front lens element that minimizes refraction at the lens-medium interface. In our experiments, we confirmed that the lenses perform in a near diffraction-limited regime over a refractive index range of 1.33 to 1.55. This gives ctASLM an axial resolving power that would be hard to match by confocal microscopy, as this would require the development of high NA lenses, which in turn would be more susceptible to refractive index mismatches.

#### Note 3 – Compabitliy of ASLM with Field Synthesis

ctASLM uses Gaussian light-sheets for excitation, which are responsible for its high-optical sectioning power. However, to optimize axial resolution and spatial duty cycle (how much of the focal plane is illuminated by the beam waist at a time), advanced light-sheets could be used. Through the recently discovered Field Synthesis theorem,^25^ such light-sheets can be generated by a simple scan of a line profile over a pupil mask. This approach can conceptually be incorporated into the presented ctASLM platform by using a normal galvanometric mirror instead of the resonant one, introducing one more telescope between the galvo mirror and the remote objective and by inserting an amplitude mask in a conjugate pupil plane (Supplementary Figure 13).

## Supplementary Figures

**Supplementary Figure 1:**
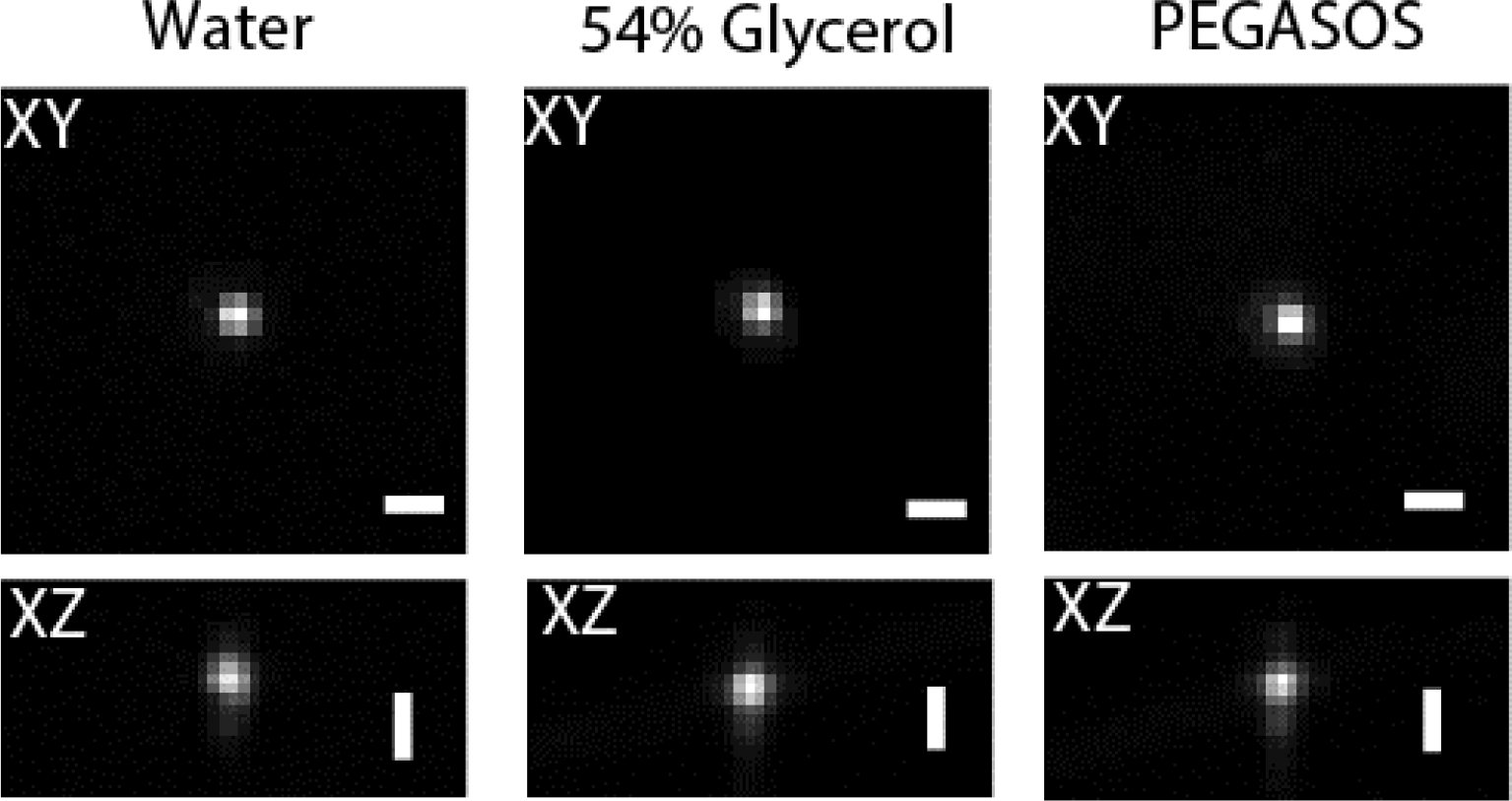
PSF of the microscope over the refractive index range of 1.33-1.54. Submicron FWHM of XY and XZ view of the PSF as shown in water (RI~1.33), 54% glycerol (RI~1.42) and PEGASOS (RI~1.543). PSF was generated by using 200 nm green fluorescent beads. Scale bar: 2um.

**Supplementary Figure 2:**
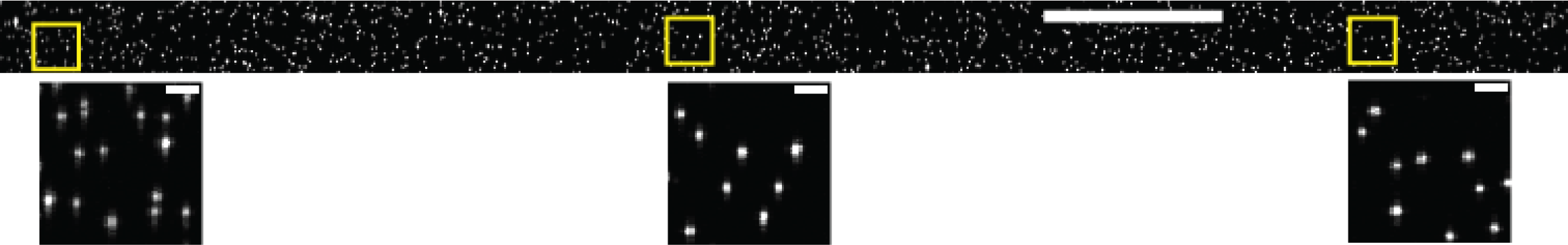
XZ view over 870×870 μm^2^ field of view depicting sub-micron isotropic resolution (0.9 ± 0.03 μm laterally and 0.85 ± 0.03 μm axially). These PSFs, generated using 200 nm green fluorescent beads embedded in a 2% agarose cube, show sub-micron light-sheet waist across the entire camera (pixel size 425 nm, 2048 × 2048× pixels). Scale bar: 100 μm and 5 μm, respectively.

**Supplementary Figure 3:**
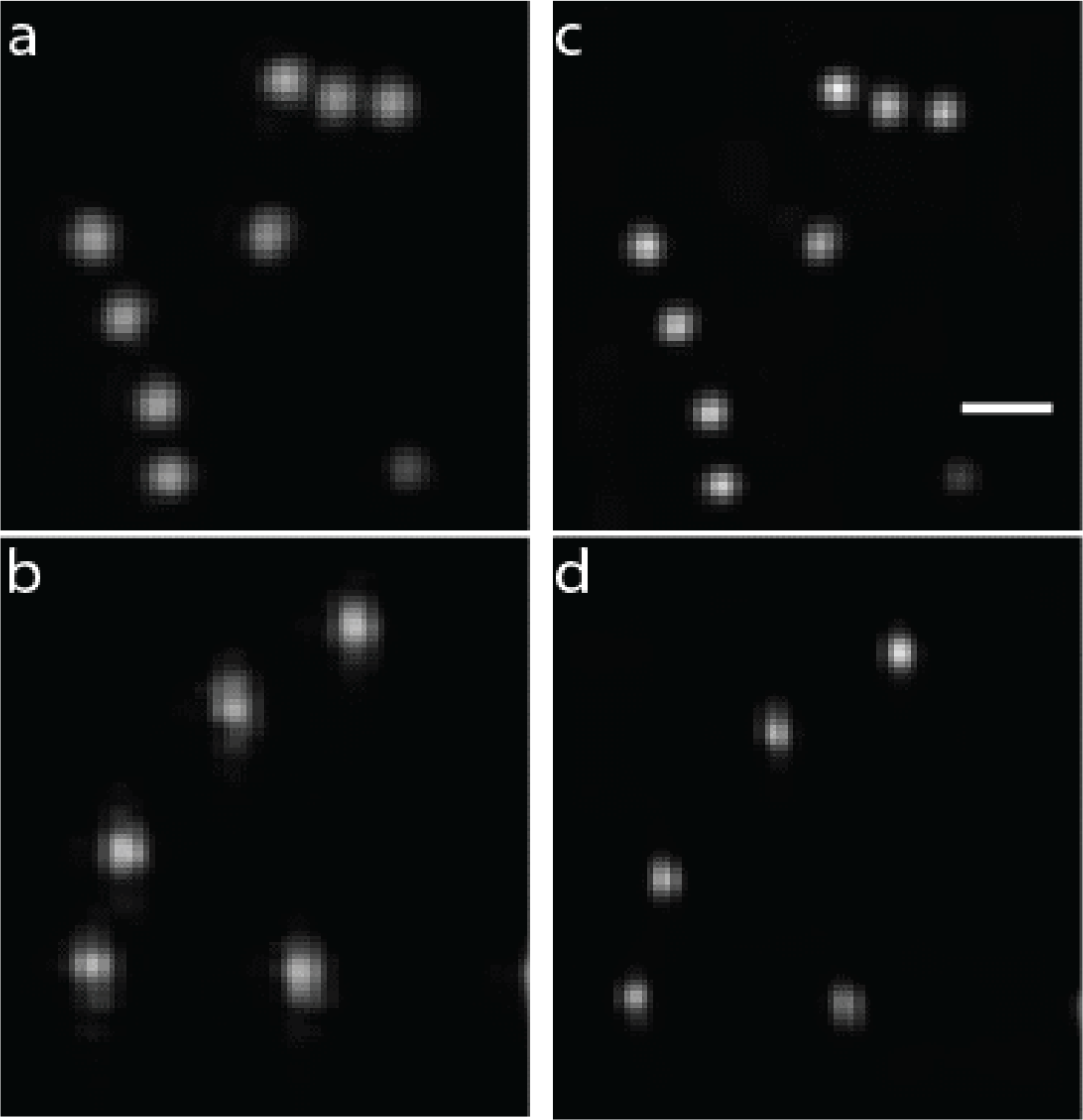
Richardson-Lucy deconvolution can improve spatial resolution by ~30%, resulting in a full-width half-maximum of fluorescent beads of 624 nm. (a,c) 200 nm fluorescent beads before and after 10 rounds of deconvolution, respectively. X-Y maximum intensity projection is shown. (b,d) axial view of fluorescent nanospheres before and after deconvolution, respectively. The data shown was resampled by a factor of two. Scale bar: 5 microns.

**Supplementary Figure 4:**
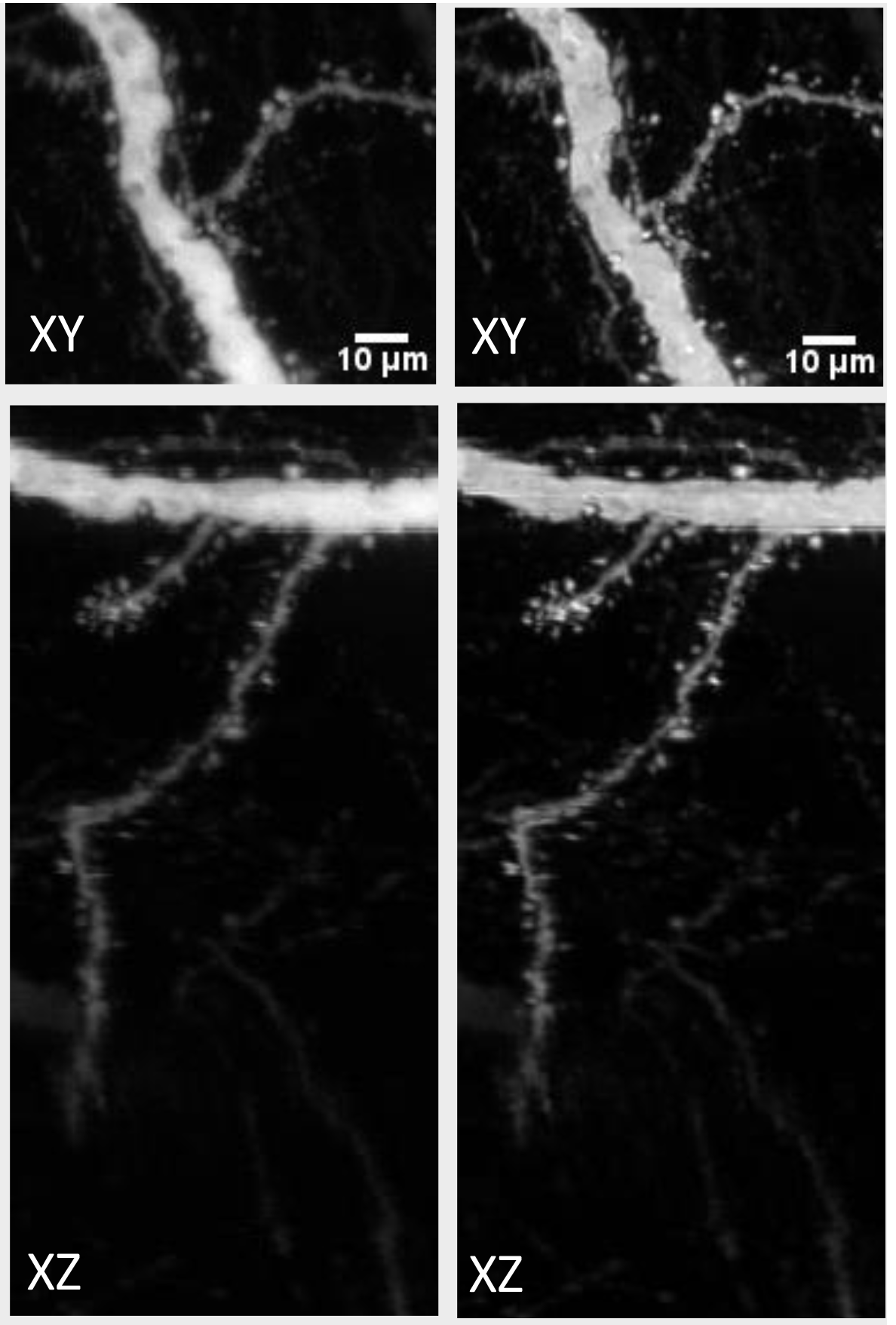
Comparison of Neurons as imaged by ctASLM before (left column) and after Richardson-Lucy deconvolution (right column), respectively. Scale bar 10 μm.

**Supplementary Figure 5:**
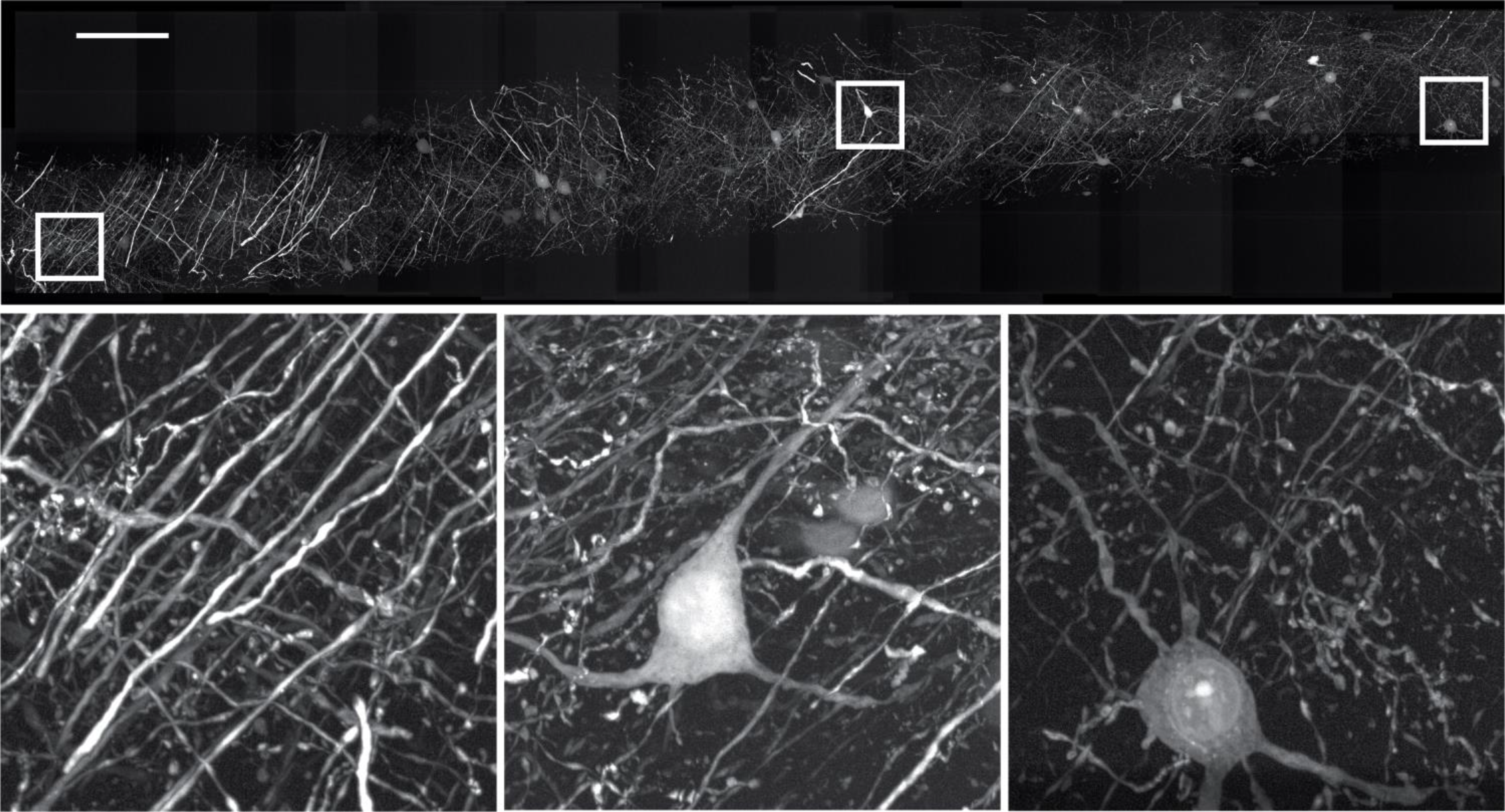
Maximum intensity projection of a chemically expanded hippocampal slice imaged with ctASLM. Insets show magnified views of the boxed regions. Scale bar 100 microns (includes magnification by expansion and by the microscope).

**Supplementary Figure 6:**
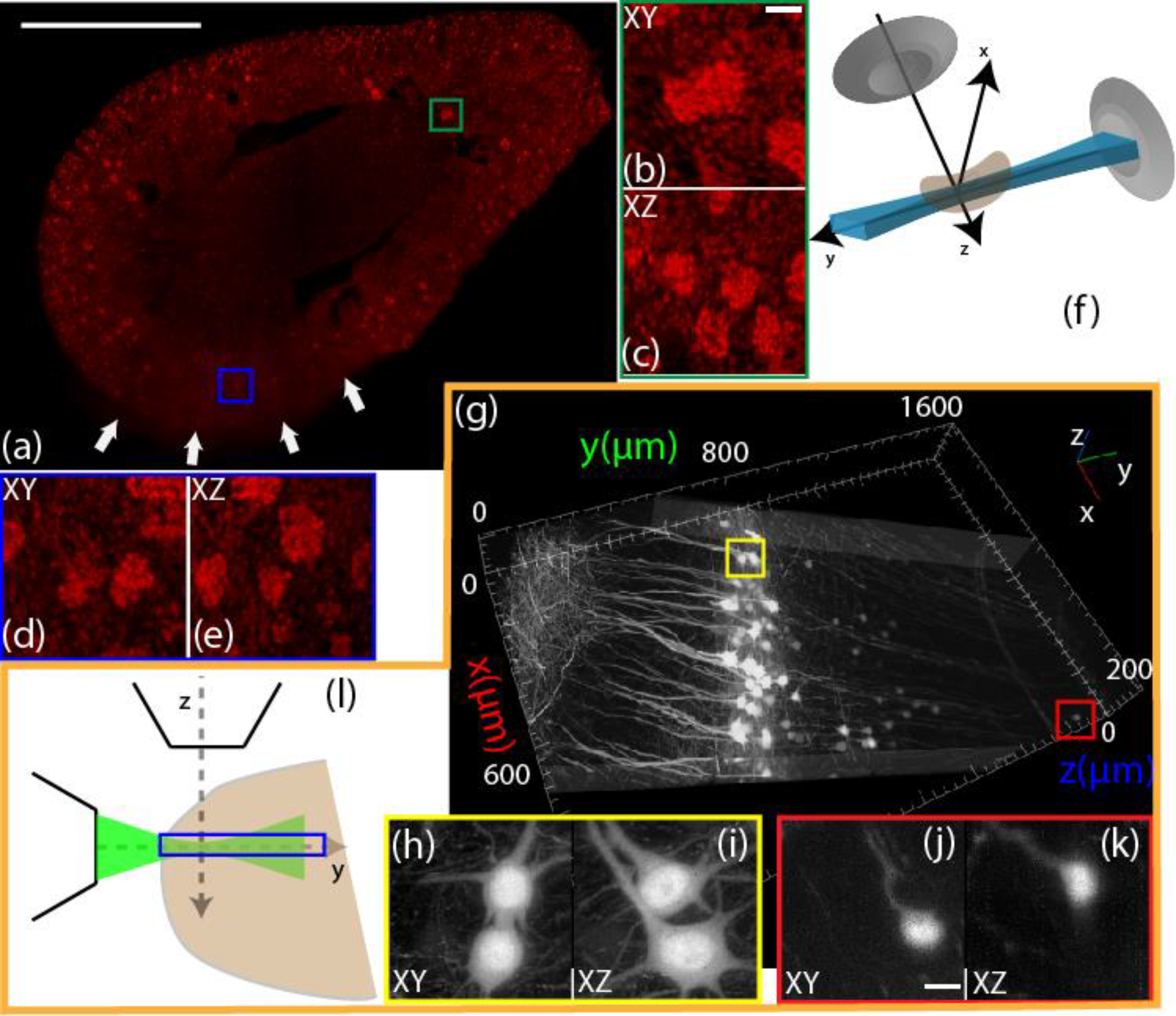
Depth performance of ctASLM within tissues. (a) mouse kidney shown in Figure 2d **and** e. White arrows point to the location where the kidney was glued to the sample holder. Scale bar 500 μm. (b-c) Magnified views of the green boxed area in (a). Scale bar 20 μm. (d-e) Magnified view of the blue boxed area in (a). (f) Schematic illustration of the illumination geometry. (g) Slice of a data volume acquired from a Clarity cleared mouse brain. (h-i) Magnified views of the boxed yellow area in (g). (j-k) Magnified view of the red boxed area in (g). (l) Location of the slice relative to the illumination and detection objective. Scale bar 20 μm.

**Supplementary Figure 7:**
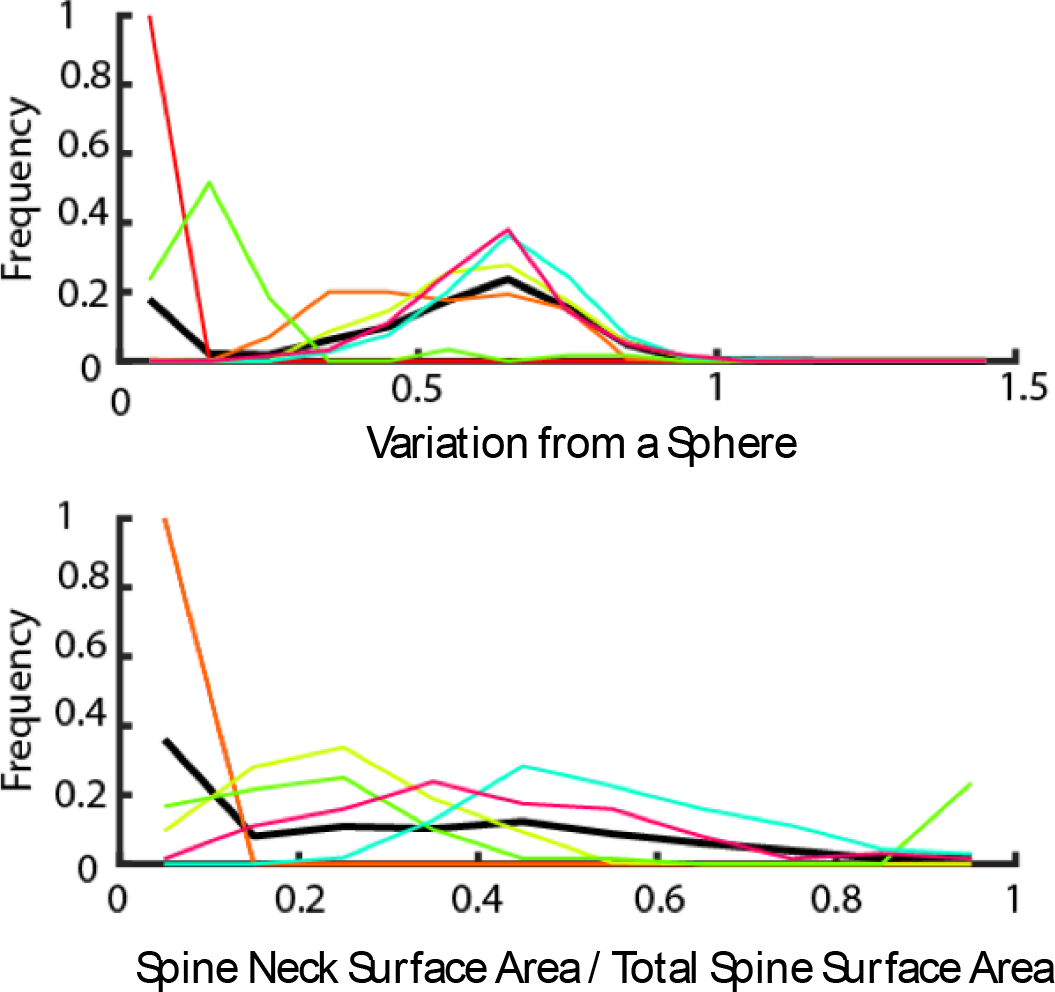
Analysis of detected spines. (Top) Variation from a sphere. (Bottom) Ratio of spine neck surface area to total spine surface area. Colors correspond to clusters identified by hierarchical clustering, and both x-axes are unitless.

**Supplementary Figure 8:**
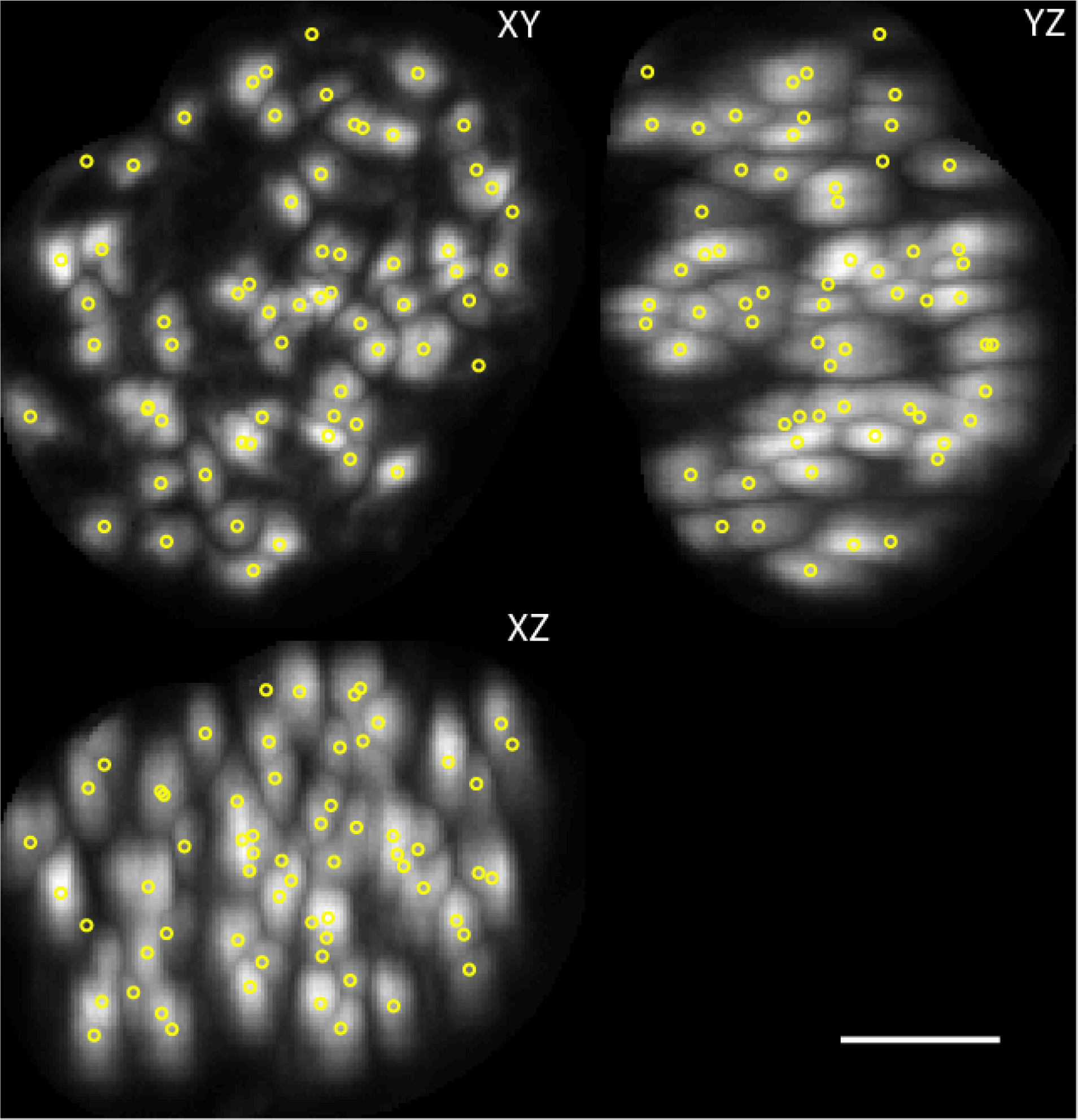
Detected endothelial cells in a gromerulli. Individual glomeruli were identified and masked using a watershed filter on data that was filtered with a 20 pixel 3D Gaussian blur. Cells within a glomerulus were then detected within the masked area using a second watershed algorithm. Scalebar 50 μm

**Supplementary Figure 9:**
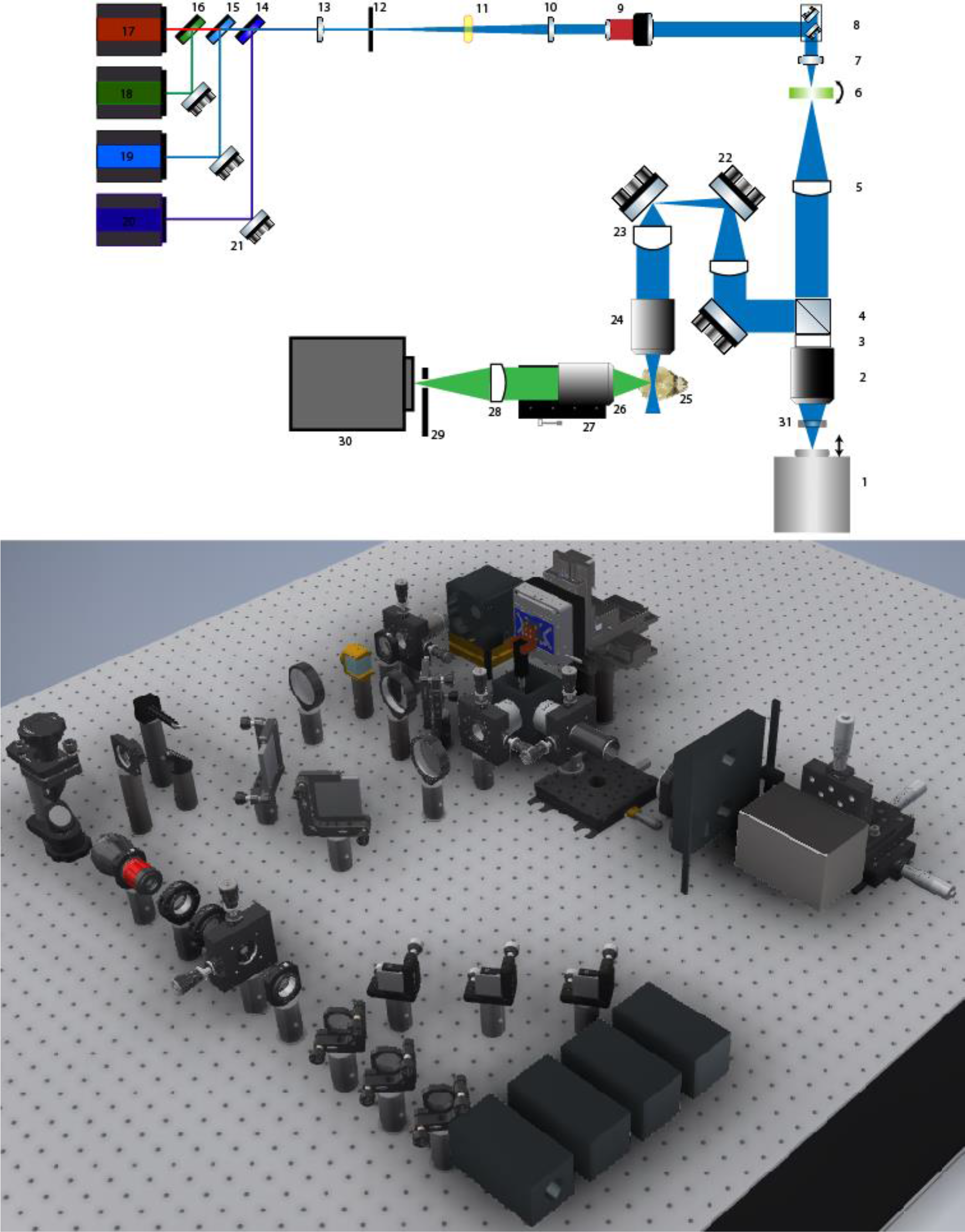
Schematic drawing (top) and CAD rendering (bottom) of the ctASLM system. A part list is provided in Supplementary Table 2.

**Supplementary Figure 10:**
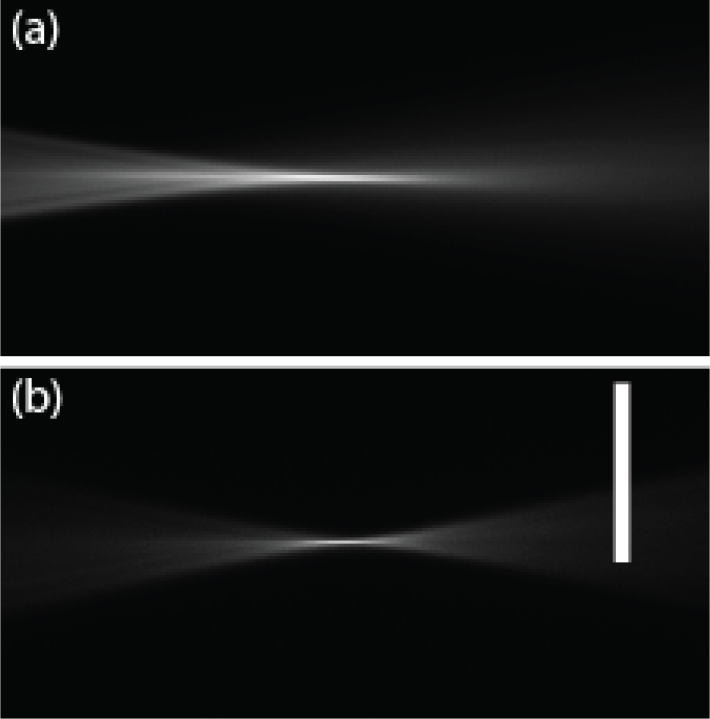
The remote focusing arm of the microscope uses an Olympus objective that is designed to image with a working distance that consists of 24 mm of air and 5 mm of water. Here, the objective was used with only an air interface, simplifying the operation of the remote focusing system. By introducing a 3 mm of glass window in the image space, spherical aberrations were eliminated. (a) and (b) show the light sheets with and without the spherical aberration by adding and removing the glass coverslip, respectively. Scale bar 50 μm.

**Supplementary Figure 11:**
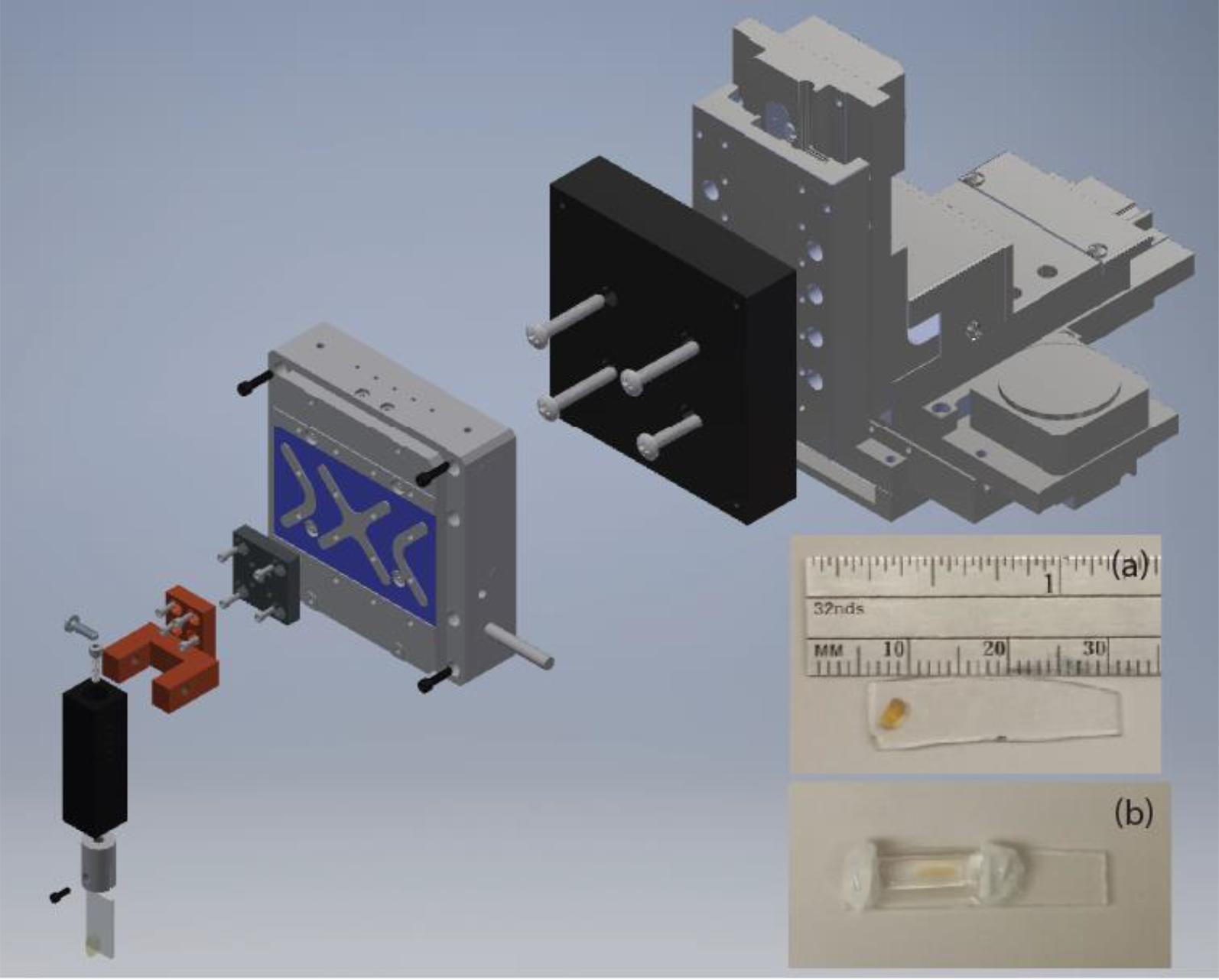
Mounting of the sample and custom sample holder assembly. Main figure: CAD rendering of the sample mount and the adapter to the sample scanning stage. (a) A neonatal mouse kidney is glued to the glass slide using superglue. (b) Bone marrow embedded in cleared agarose. The agarose-cylinder is then held to the glass slide using silicone form the two ends.

**Supplementary Figure 12:**
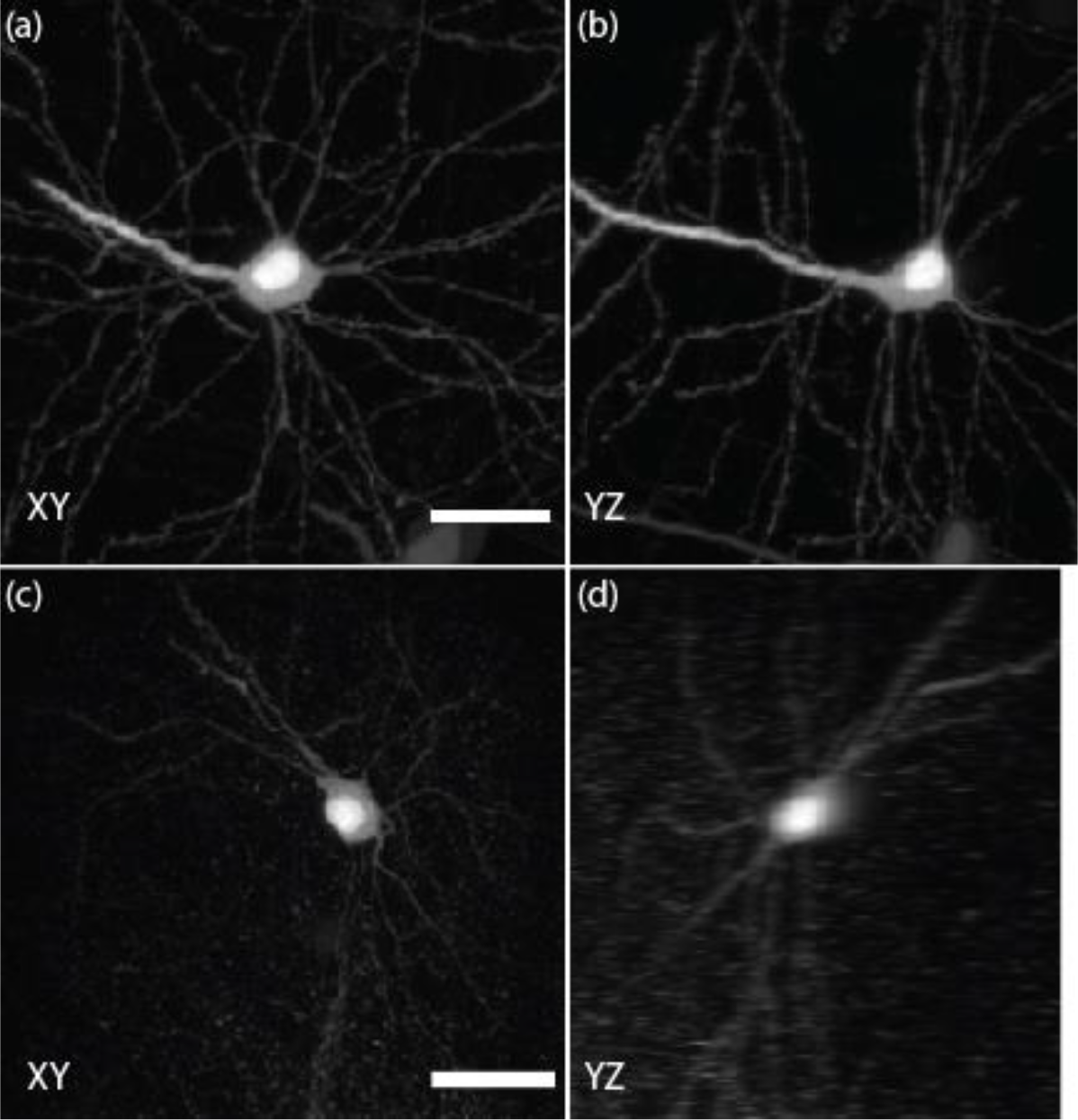
Comparison of ctASLM’s imaging performance to that of confocal microscope. Cortex of a PEGASOS cleared mouse brain expressing THy1-GFP (a) XY and (b) YZ were acquired using ctASLM. (c) and (d) are a representative area imaged using a confocal microscope. Scale bar: 25 μm.

**Supplementary Figure 13:**
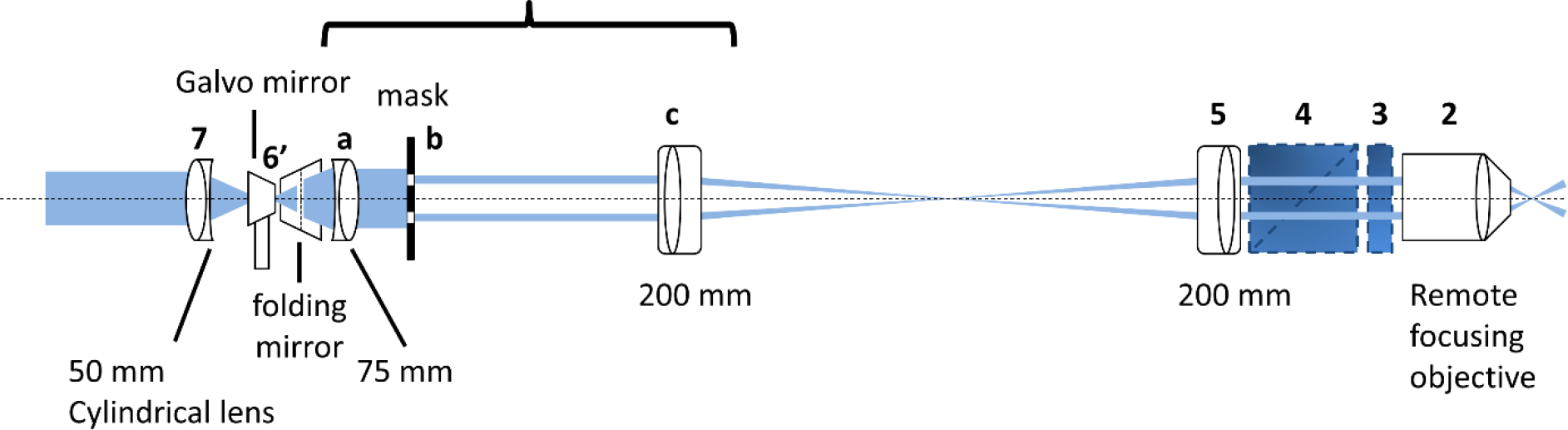
Field Synthesis unit. The main components of Field synthesis consist of a galvo mirror (**6’**) and a mask (**b**). By inserting components **a**, **b**, and **c** into the optical path shown in Supplementary Figure 9, Field Synthesis can be implemented into ctASLM. Components **2**,**3**,**4**,**5**, and **7** can remain the same. The mask (**b**) is placed in the conjugated plane of the back focal plane of the remote focusing objective.

### Supplementary Tables

**Supplementary Table 1:** Lateral and axial resolution of ctASLM in different clearing media. 200 nm fluorescent nanospheres were used for PSF measurements. Five beads were measured for each media.

|  | Water (RI~1.33) | Glycerol (RI~1.42) | PEGASOS (RI~1.54) |
| --- | --- | --- | --- |
| $\delta x$ ($\mu\text{m}$) | 0.90±0.03 | 0.90±0.05 | 0.98±0.03 |
| $\delta y$ ($\mu\text{m}$) | 0.90±0.02 | 0.97±0.02 | 0.92±0.04 |
| $\delta z$ ($\mu\text{m}$) | 0.85±0.03 | 0.92±0.04 | 0.94±0.02 |

**Supplementary Table 2:**
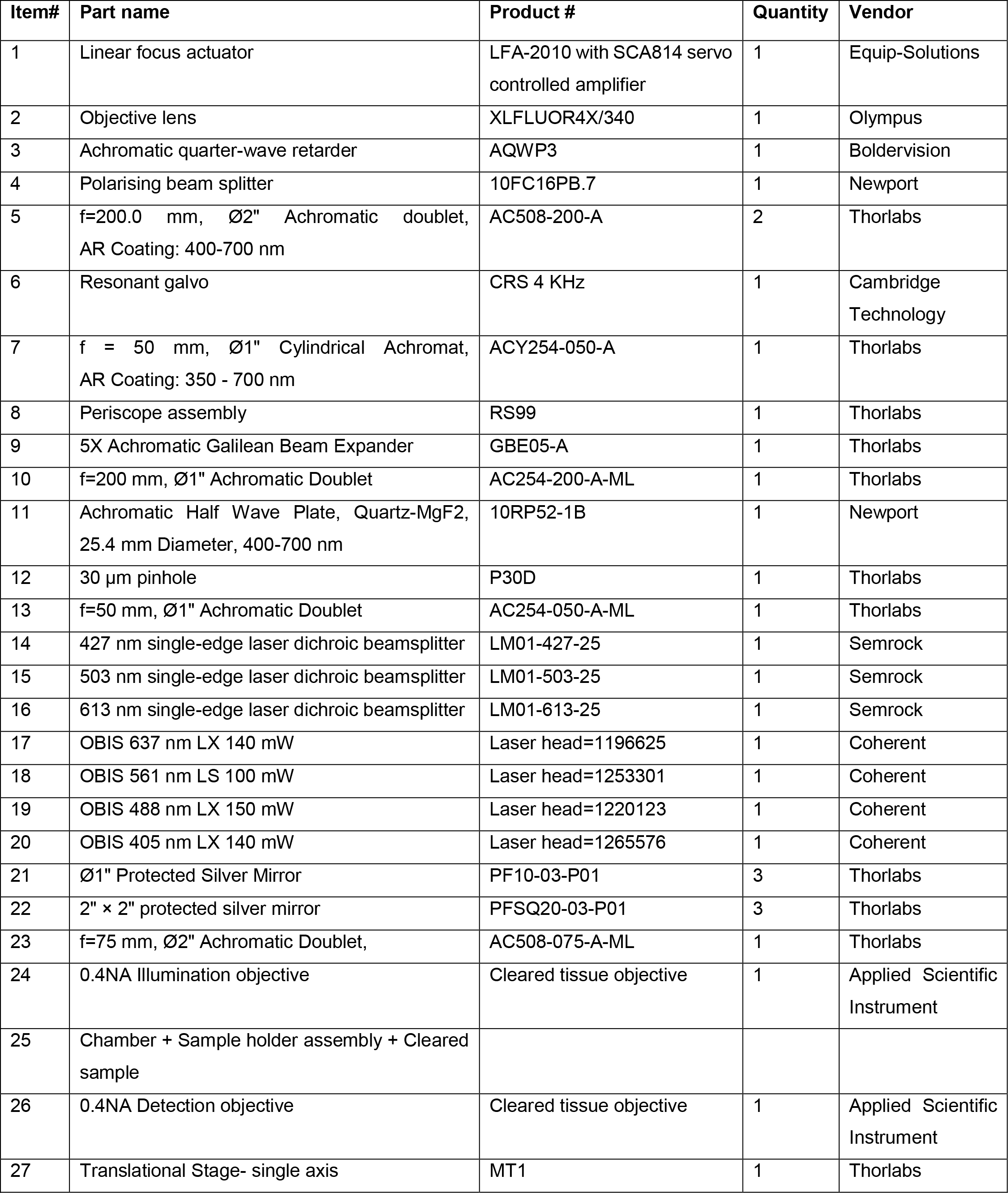
Parts list for the ctASLM microscope.

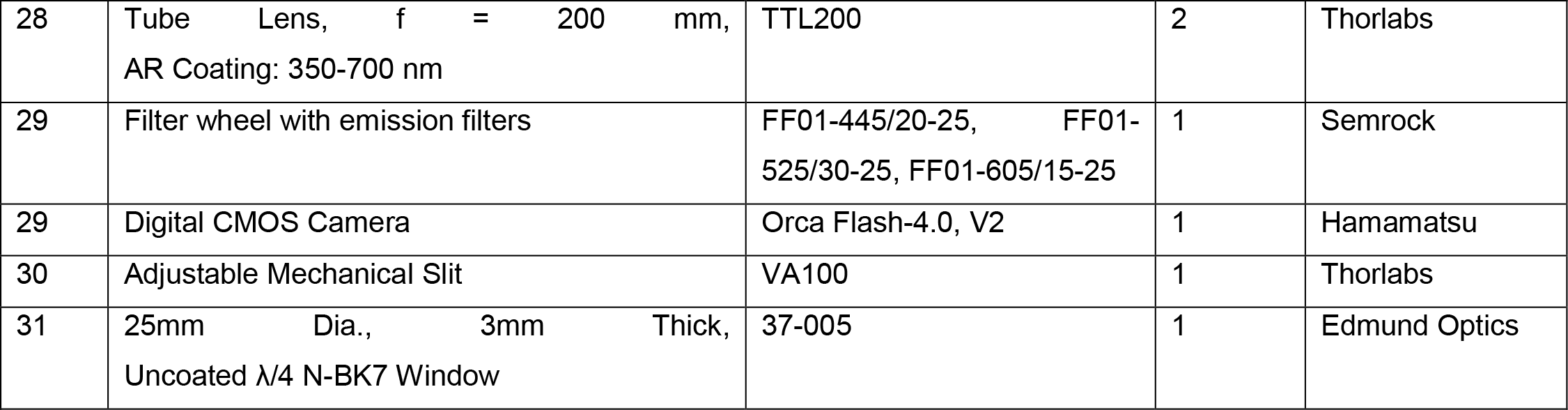

### Supplementary Movies

**Supplementary Movie 1.**
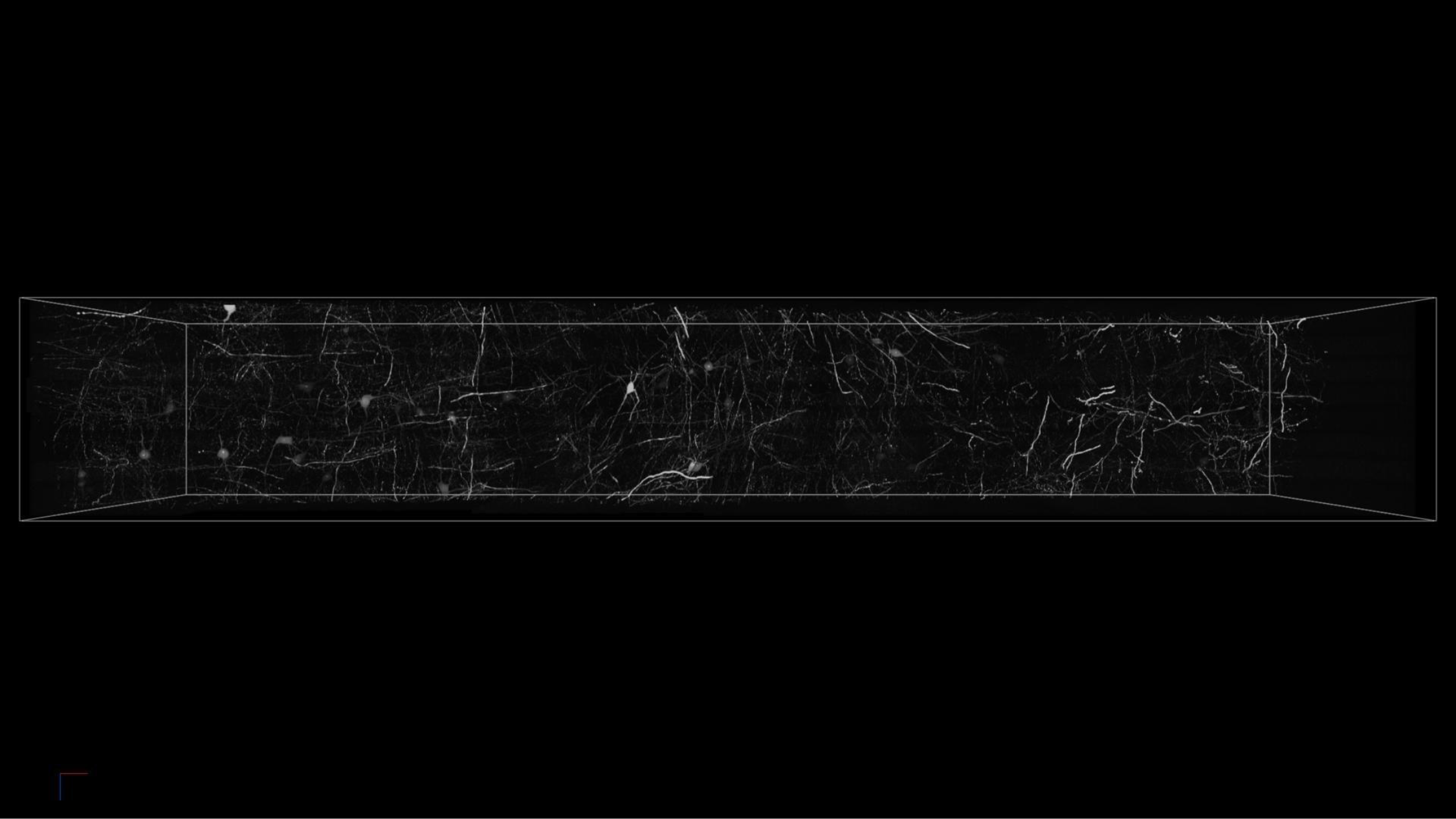
Expansion Microsocopy. Chemically expanded hippocampal slice with Thy1-GFP labeled neurons as imaged with ctASLM. Image dimensions are 8137 × 1645 × 3020 voxels, with a voxel size in sample space (due to tissue expansion) of 100 nm. Thus, the sample dimensions are 823 × 166 × 302 μm^3^.

**Supplementary Movie 2:**
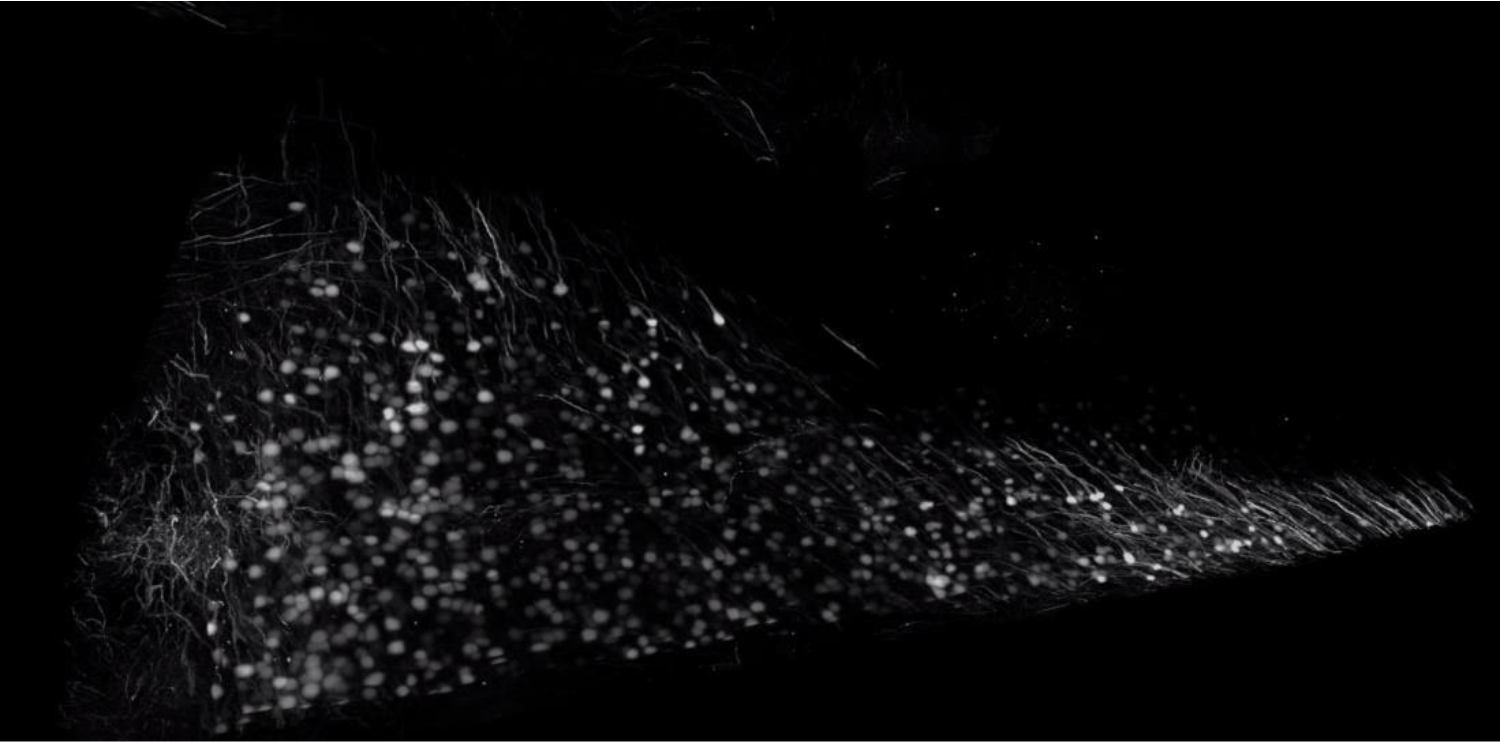
A 5 × 3 × 3 mm^3^ section of Thy1-eYFP-expressing neurons from the cortical section an adult mouse brain cleared using CLARITY. A 54% glycerol solution (RI~1.42) was used for refractive index matching and imaging.

**Supplementary Movie 3:**
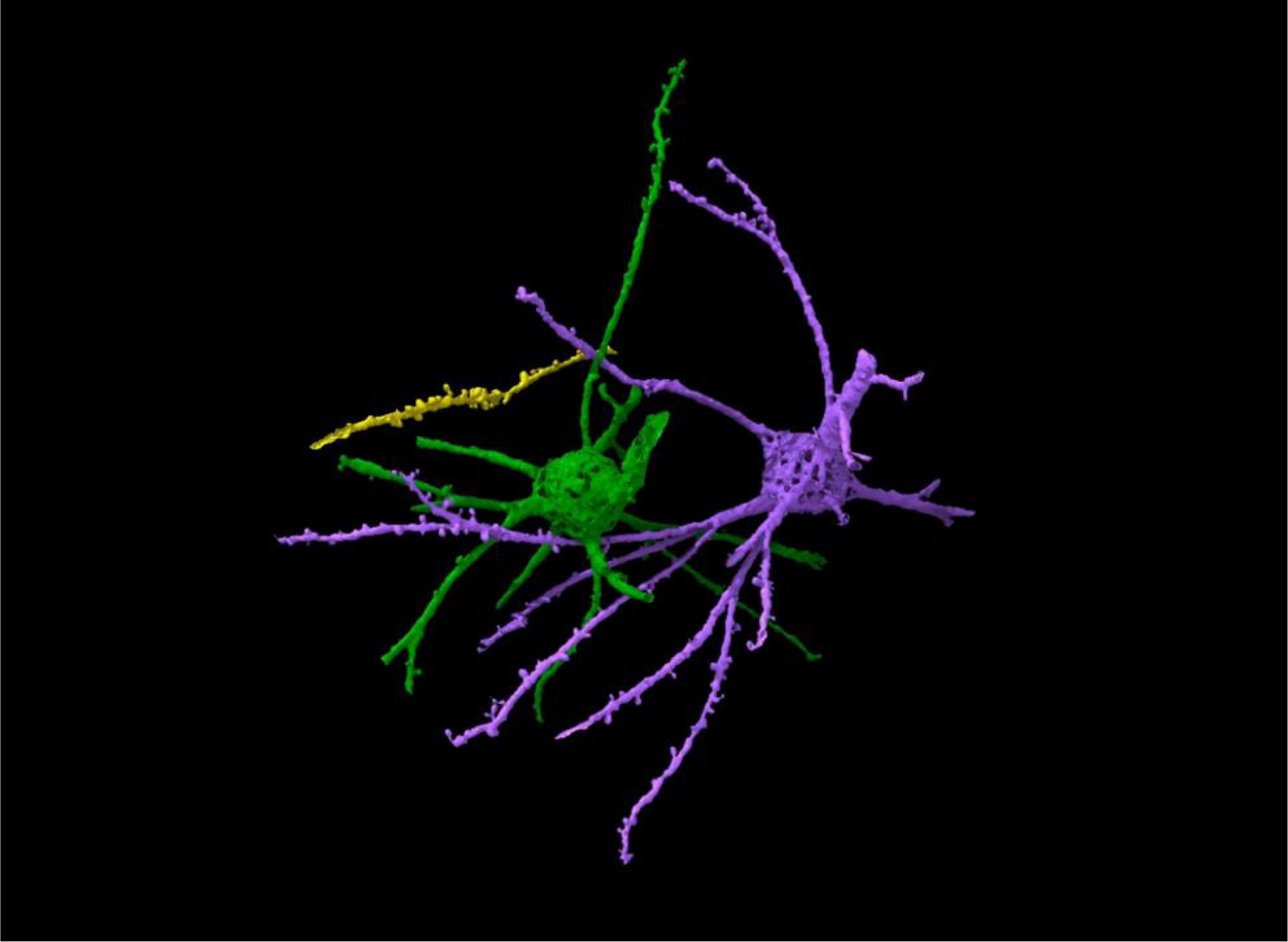
Segmented Thy1-GFP cortical neurons from a CLARTY cleared mouse brain, in a sub-region spanning 170 × 170 × 60 μm^3^.

**Supplementary Movie 4:**
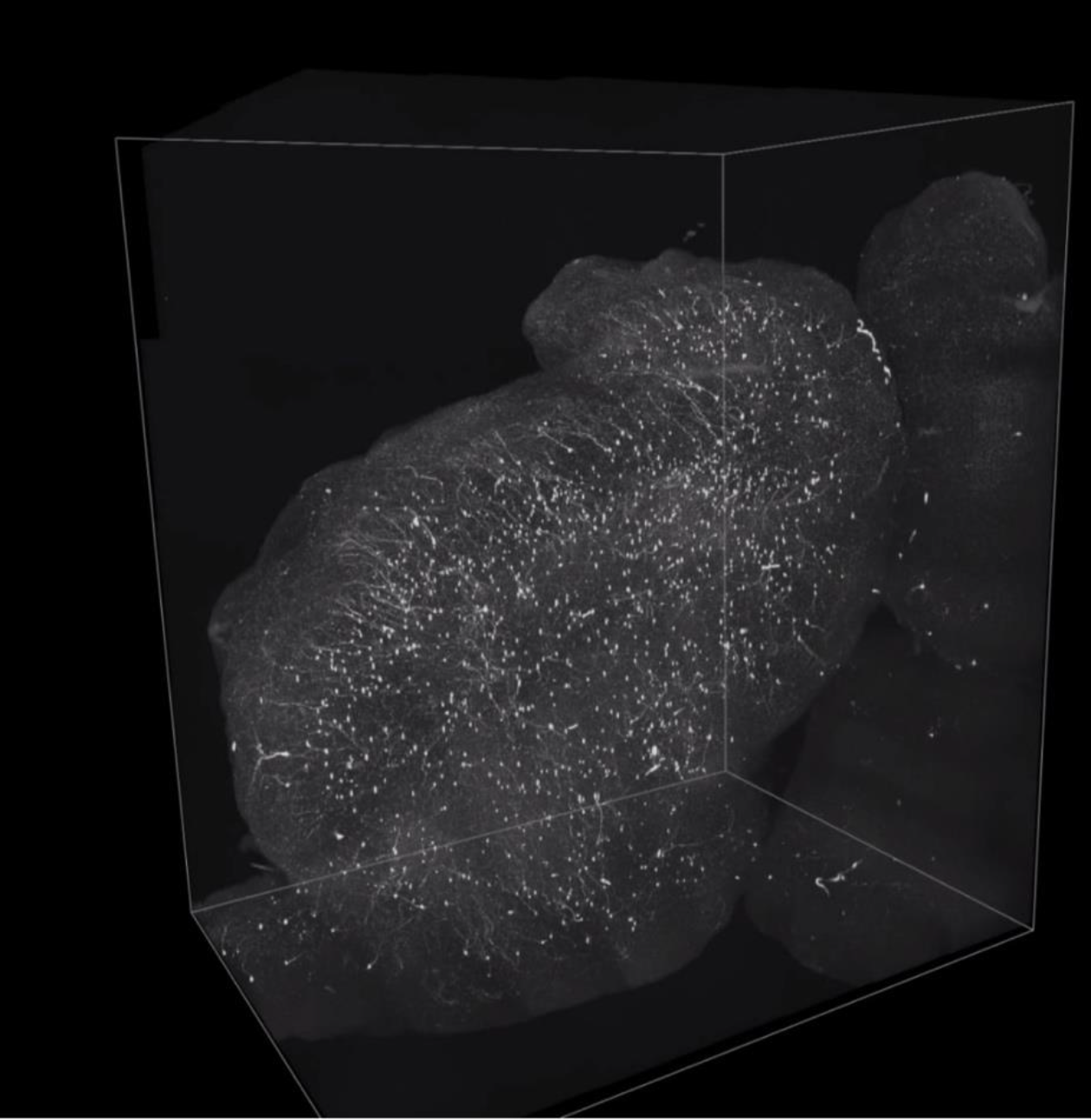
Olfactory bulb neurons labeled with a stimulus-inducible TdTomato reporter, and cleared with PEGASOS. Imaging region spans 2.8 × 2 × 0.66 mm^3^.

**Supplementary Movie 5:**
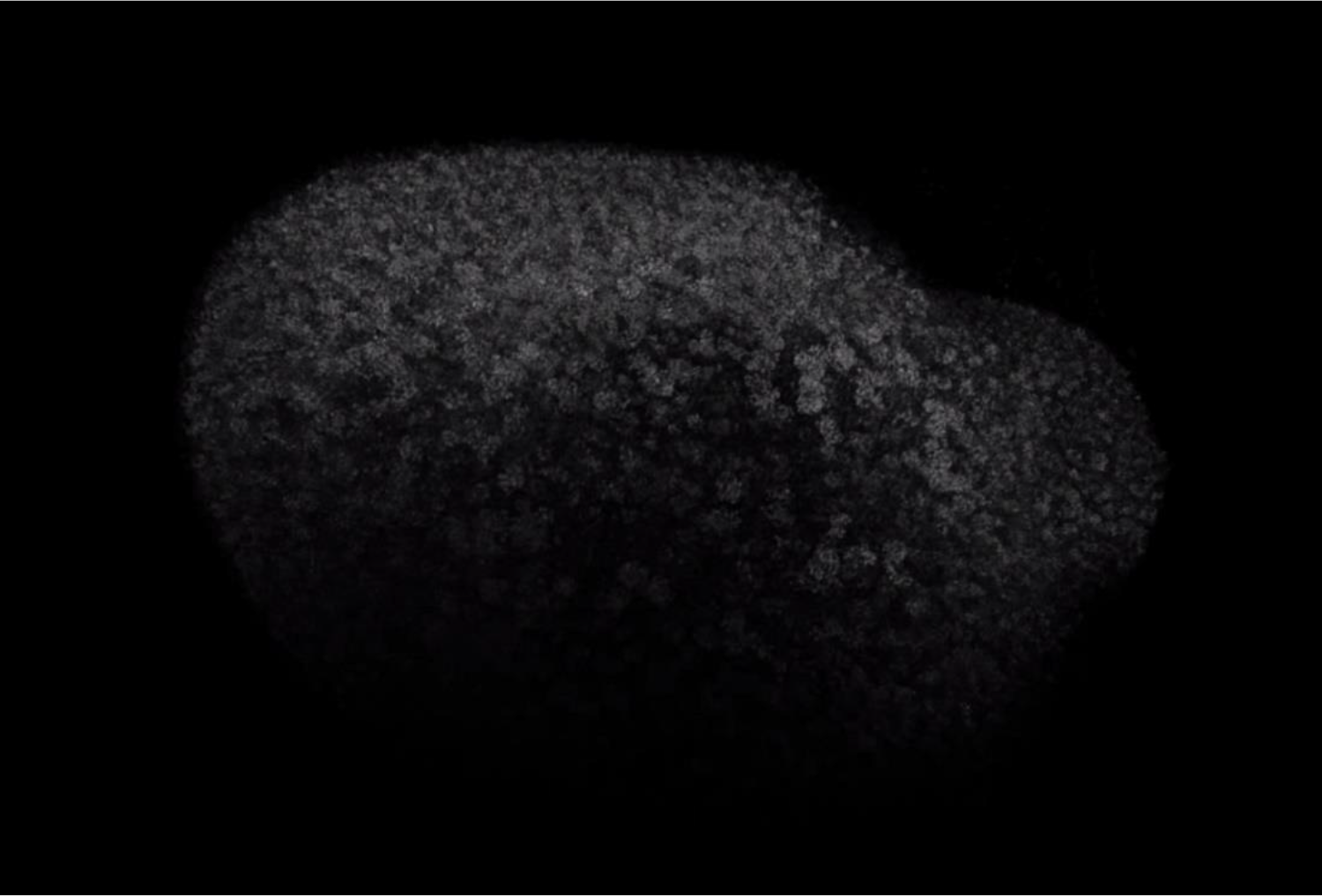
BABB cleared mouse kidney labeled with Flk1-GFP in an imaging volume of 3 × 2.3 × 2.3 mm^3^.

**Supplementary movie 6:**
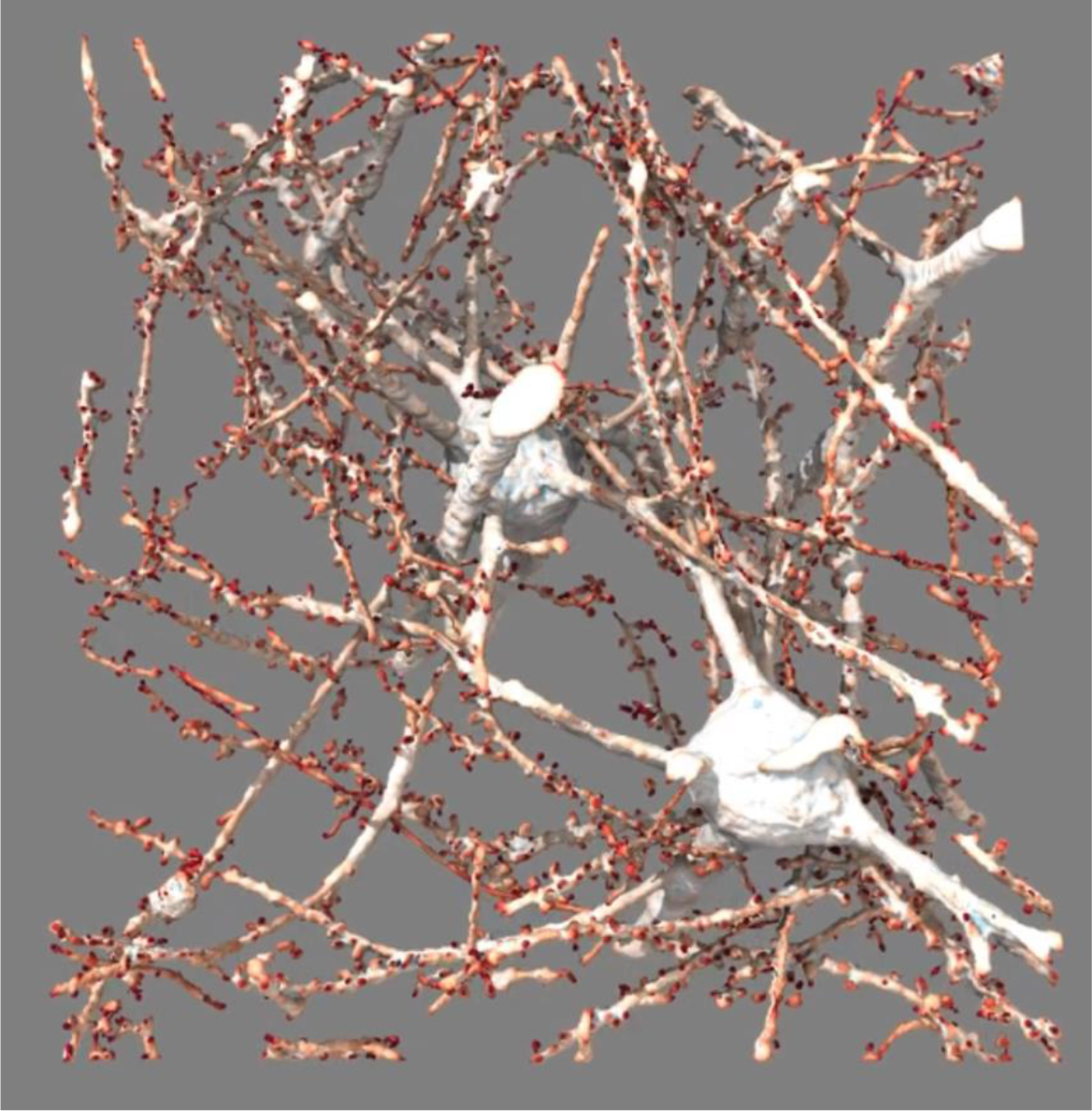
Detection of spines on Thy1-EYFP neurons based on curvature over a 176 × 190 × 60 μm^3^ volume.

**Supplementary Movie 7:**
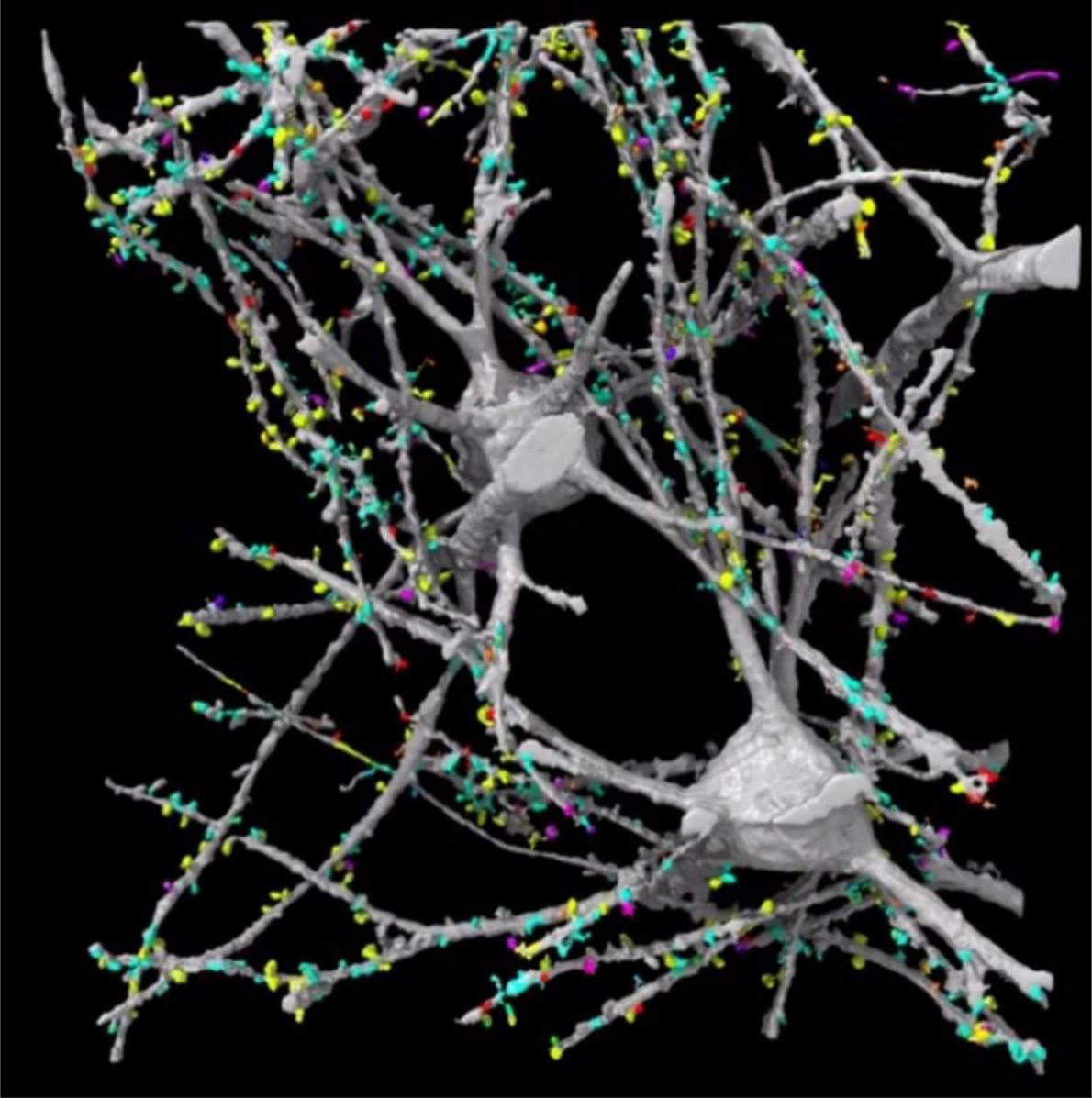
Clustering of spines following principal component analysis of their morphological parameters.

**Supplementary Movie 8:**
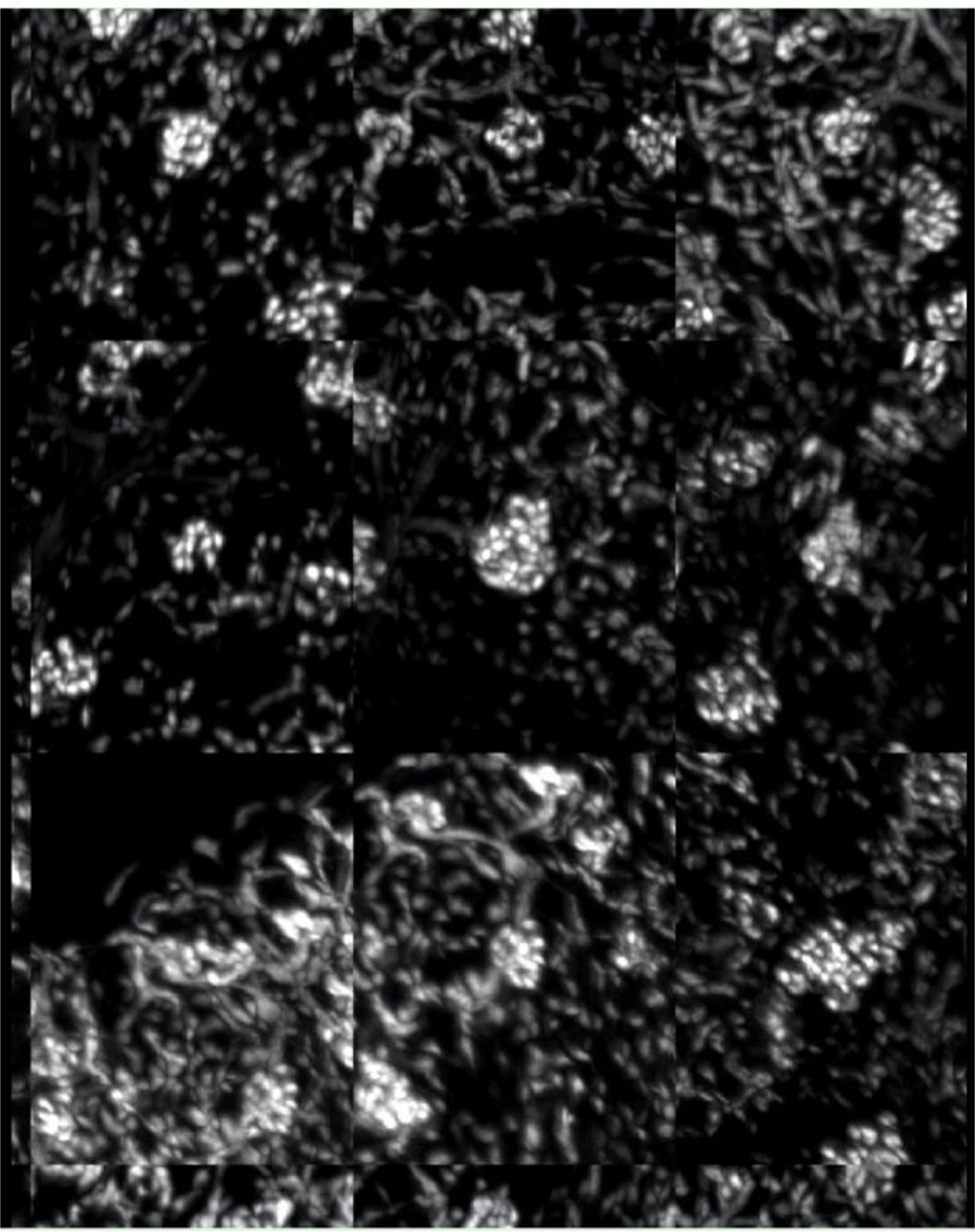
Automatically detected glomeruli, assembled into a mosaic. Each glomerulus is located at the center of each sub-image.

**Supplementary Movie 9:**
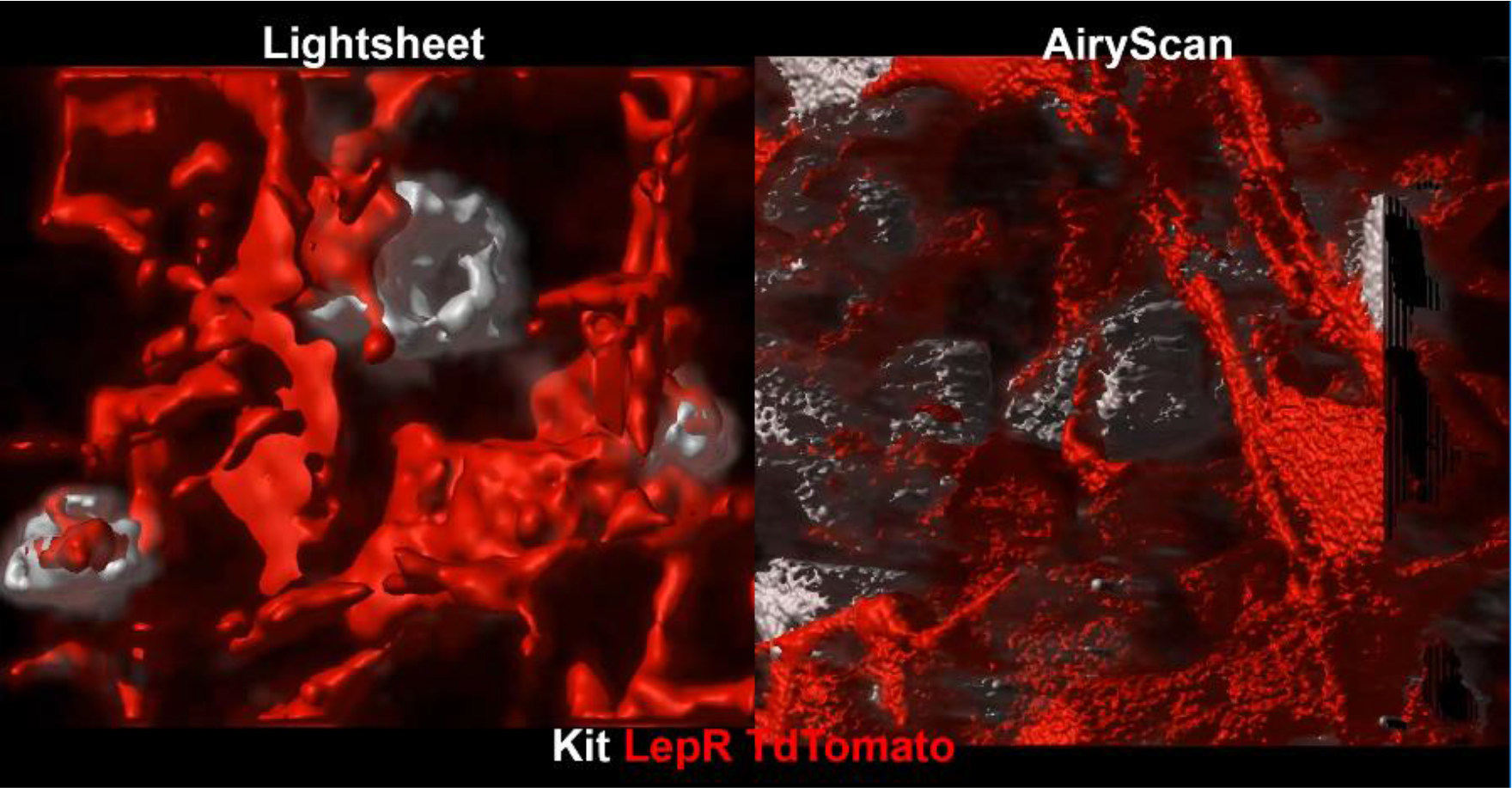
Imaging of an hematopoetic stem cell niche using ctASLM (left) and AiryScan (right).

